# Telomere-to-telomere genome assembly of a male goat reveals novel variants associated with cashmere traits

**DOI:** 10.1101/2024.03.03.582909

**Authors:** Hui Wu, Ling-Yun Luo, Ya-Hui Zhang, Chong-Yan Zhang, Jia-Hui Huang, Dong-Xin Mo, Li-Ming Zhao, Zhi-Xin Wang, Yi-Chuan Wang, He-Hua EEr, Wen-Lin Bai, Di Han, Xing-Tang Dou, Yan-Ling Ren, Renqing Dingkao, Hai-Liang Chen, Yong Ye, Hai-Dong Du, Zhan-Qiang Zhao, Xi-Jun Wang, Shan-Gang Jia, Zhi-Hong Liu, Meng-Hua Li

**Affiliations:** College of Animal Science and Technology, China Agricultural University, Beijing, China; Northern Agriculture and Animal Husbandry Technical Innovation Center, Chinese Academy of Agricultural Sciences, Hohhot, China; College of Animal Science, Inner Mongolia Agricultural University, Hohhot, China; Laboratory of Herbage Improvement and Grassland Agro-ecosystems; Key Laboratory of Grassland Livestock Industry Innovation, Ministry of Agriculture and Rural Affairs; Engineering Research Center of Grassland Industry, Ministry of Education; College of Pastoral Agriculture Science and Technology, Lanzhou University, Lanzhou, China; Institute of Animal Science, NingXia Academy of Agriculture and Forestry Sciences, Yinchuan, China; College of Animal Science and Veterinary Medicine, Shenyang Agricultural University, Shenyang, China; Modern Agricultural Production Base Construction Engineering Center of Liaoning Province, Liaoyang, China; Liaoning Province Liaoning Cashmere Goat Original Breeding Farm Co., Ltd., Liaoyang, China; Shandong Binzhou Academy of Animal Science and Veterinary Medicine, Binzhou, China; Gannan Institute of Animal Husbandry Science, Hezuo, China; Beijing Lvyeqingchuan Zoo Co., Ltd., Beijing, China; Zhongwei Goat Breeding Center of Ningxia Province, Zhongwei, China; Jiaxiang Animal Husbandry and Veterinary Development Center, Jining, China; College of Grassland Science and Technology, China Agricultural University, Beijing, China

**Keywords:** telomere-to-telomere assembly, goat, acrocentric chromosome, Y chromosome, gap-free, cashmere, domestication

## Abstract

A complete goat (*Capra hircus*) reference genome enhances analyses of genetic variation, thus providing novel insights into domestication and selection in goats and related species. Here, we assembled a telomere-to-telomere (T2T) gap-free genome (2.86 Gb) from a cashmere goat (*T2T-goat1.0*), which included a Y chromosome of 20.9 Mb. With a base accuracy of >99.999%, *T2T-goat1.0* corrected numerous genome-wide structural and base errors in previous assemblies and added 288.5 Mb of previously unresolved regions (PURs) and 446 newly assembled genes to the reference genome. We sequenced the genomes of five representative goat breeds for PacBio reads, and used *T2T-goat1.0* as a reference to identify a total of 63,417 structural variations (SVs) with up to 4711 SVs (7.41%) in the PURs and 507 overlapping genes significantly enriched (*P*_adj._ < 0.05) in the keratin filament pathway. *T2T-goat1.0* was applied in population analyses of global wild and domestic goats, which revealed 32,419 SVs and 25,397,794 SNPs, including 870 SVs and 545,026 SNPs in the PURs. Also, our analyses revealed a set of novel selective variants and genes associated with domestication (e.g., *NKG2D* and *ABCC4*) and cashmere traits (e.g., *ABCC4* and *ASIP*).

## Introduction

Goats (*Capra hircus*) were domesticated in the Fertile Crescent *ca*. 10,500 years before present (BP)^1^. Footprints of natural and artificial forces have shaped their genomes through the accumulation of genomic variants. A complete reference genome is crucial for improving the analysis of genomic variants involved in the genetics, selection and evolution of goats and related species^2-4^. With the advancement of sequencing technologies, several goat genome assemblies, for example *CHIR_1.0*, have been released, followed by updates to *CHIR_2.0*^5^ (GenBank accession no. GCA_000317765.2), *Saanen_v1*^6^ (GCA_015443085.1), and *ARS1.2*^7^ (GCF_001704415.2). Nevertheless, in addition to unplaced contigs, the assemblies have unfilled gaps (e.g., 169 gaps in the most improved reference *Saanen_v1* of a Sannen goat^6^) in the pseudochromosomes, many of which might cover the gene bodies and cause a loss of uncovered genes. Additionally, the complete sequences of many centromeres, telomeres, repetitive regions and Y chromosome have not been found in these published goat reference genomes, which limits our knowledge of their organization, structure, evolution, and function^8,9^.

Advances in long-read sequencing technology, especially ultralong ONT reads, have greatly facilitated the achievement of telomere-to-telomere (T2T) genome assemblies without any gaps, including those of human^10,11^, Arabidopsis^12^, watermelon^13^, maize^14^, soybean^15^, rice^16^, and chicken^17^. Complex genome regions of autosomes and chromosomes X and Y unlocked in the T2T reference genome of human *T2T-CHM13* have revealed quite a few novel and rare variants and genes^18,19^, providing new insights into their maternal and paternal evolutionary patterns^20^. In goats, except for the metacentric Y chromosome, all the autosomes and the X chromosome are acrocentric, which impedes the complete assembly of centromeres and telomeres^6,7^. Acrocentric chromosomes are typically involved in the frequent genomic recombination^21^; thus, variations in the unresolved regions of acrocentric chromosomes are the key to understanding chromosomal evolution, genome rearrangement and selective signatures in goats and related species^22^.

To date, T2T reference genomes of large vertebrates, in addition to those of humans, have been rarely reported with all complete chromosomes^11^. Here, we described the T2T gap-free genome assembly of the cashmere goat (*T2T-goat1.0*), which includes all autosomes and chromosomes X and Y, using ultralong ONT and PacBio HiFi reads together with Hi-C and Bionano data. We revealed the genomic organization, variations, and genes in previously unresolved regions, mostly for centromeres, telomeres, and the Y chromosome. We showed the advantages of using the *T2T-goat1.0* genome as a reference for population genomics analyses, such as sequence alignment, variation calling, genetic structure and selective signal detection.

## Results

### T2T genome assembly

We selected a buck from the Inner Mongolia cashmere goat, a representative indigenous goat breed in China (Fig. 1a), for the T2T genome assembly. We generated a total of 328.44 Gb of ultralong ONT (114.8× coverage), 141.03 Gb of PacBio HiFi (49.3× coverage), 435.76 Gb of Hi-C data (152.2× coverage), and 1188.9 Gb of Bionano optical mapping data (Supplementary Table 1). The initial goat genome assembly (GV1) was built based on PacBio HiFi data, and 97 contigs were scaffolded as the GV2 assembly with Bionano data. Using Hi-C data, the contigs/scaffolds were anchored and oriented onto 30 pseudochromosomes corresponding to 29 autosomes and the X chromosome of the GV3 assembly (Supplementary Fig. 1), with 145 gaps and an average of ∼4.8 gaps per pseudochromosome (Supplementary Fig. 2). Our assembly results showed that all the autosomes and the X chromosome were acrocentric (Fig. 1b), causing great challenges in the complete chromosomal end assembly. For example, a tangle of centromeric regions among four chromosomes, CHI12, CHI13, CHI19 and CHI22, was shown in the assembly graph string (Supplementary Fig. 3). Therefore, the gaps were enriched in centromeric regions (Supplementary Fig. 2a), with the longest gap (∼4 Mb) located on the centromeric region of CHI14 (*C. hircus* chromosome 14), which resulted from repetitive sequences. We employed ultralong ONT reads to fill these gaps via extension or local assembly, depending on the gap size and complexity. The filled gaps were confirmed for reliability based on the alignment of the HiFi and ONT long reads, and the genome GV4 was created accordingly (Fig. 1b and Supplementary Fig. 2b). To construct the telomeric regions accurately, HiFi reads containing telomeric repeat sequences were used to independently assemble and replace the sequence errors at the ends of all chromosomes.

**Fig. 1.**
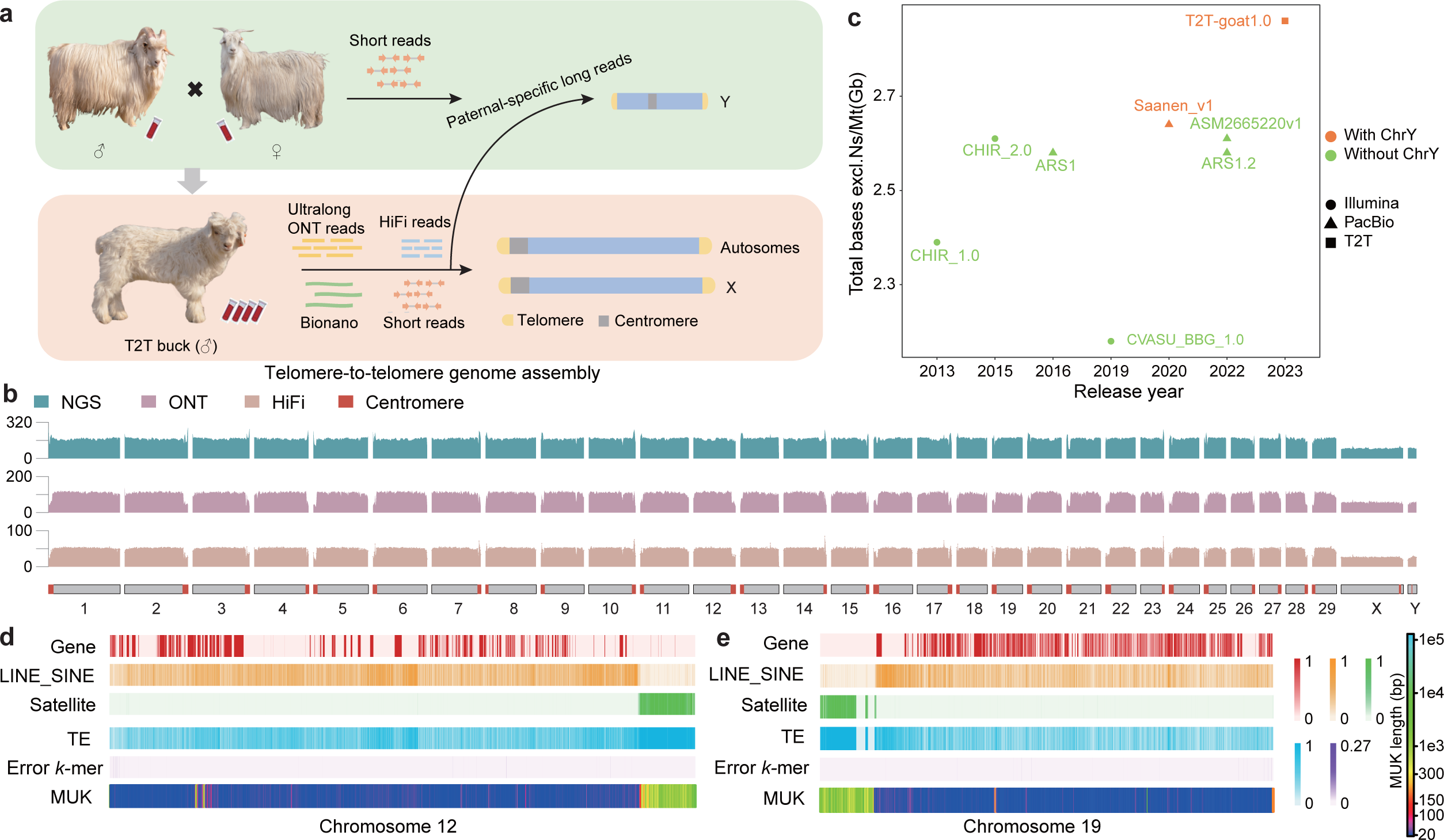
A complete goat genome with 29 autosomes and chromosomes X and Y. **a,** Genome assembly strategy for *T2T-goat1.0* of a buck. Independent Y chromosome assembly was performed with assistance of MGI short reads from the buck and its parents. **b,** Whole-genome coverage of mapped short (NGS for MGI) and long reads (HiFi and ONT) along all 31 chromosomes of *T2T-goat1.0* in 1-Mb windows. The red color represents the centromeric region on each chromosome. **c,** Genome size distribution of the available seven chromosome-level goat assemblies in a comparison to that of *T2T-goat1.0*, *CHIR_1.0* (GCF_000317765.1), CHIR_2.0 (GCA_000317765.2), *ARS1* (GCF_001704415.1), CVASU_BBG_1.0 (GCA_004361675.1), *Saaven_v1* (GCA_015443085.1), *ARS1.2* (GCF_001704415.2), *ASM2665220v1* (GCA_026652205.1). Circles represent assemblies generated using Illumina, with triangles for the ones using PacBio and squares for *T2T-goat1.0*. The orange color denotes an assembly of a buck with a chromosome-level Y chromosome, and the green color represents a doe without Y chromosome assembled. Ns and Mt are for the gaps and mitochondria respectively. **d, e,** Genome features of chromosomes 12 and 19. The following information is provided from top to bottom: the gene density, the density of LINEs and SINEs, the satellite density, the TE density, and the minimum unique *k*-mer (MUK) per 100 kb. All the features are shown in 10-kb windows, except for MUK.

Based on next-generation sequencing (NGS) data from the assembled buck and its parents (Fig. 1a), paternal-specific *k*-mers were generated and used to pick ONT reads of paternal origin for Y-chromosome assembly. Y chromosomal contigs *de novo* assembled from HiFi reads were used to improve the Y chromosome assembly based on paternal ONT reads, including scaffolding, filling gaps and correction. As a result, we generated a complete Y chromosome with two telomeres spanning 20.9 Mb.

### Validation of the T2T goat genome assembly

All the chromosomes in GV4 were polished using our local pipeline based on the ONT and PacBio HiFi long reads and the NGS short reads. The final T2T genome assembly, *T2T-goat1.0*, covering all the 29 autosomes and X and Y chromosomes, was obtained with a total length of 2.86 Gb (Table 1 and Fig. 1b). This size was slightly longer than the estimated genome size of 2.76 Gb obtained using the NGS short reads. Multiple strategies were further used to assess the integrity and accuracy of *T2T-goat1.0*. The average consensus quality value (QV) was estimated to be 54.18, with an increase from ∼35 in the assembly GV4, indicating a base accuracy >99.999%. A completeness of 95.71% was obtained, and estimates of QV ranged from 50.07 to 59.31 across all the chromosomes, with 49.10 for the Y chromosome (Supplementary Table 2). The primary data, including HiFi and ONT long reads and short reads, were all mapped to *T2T-goat1.0*, and the read coverage was found to be evenly distributed across nearly all the chromosomes (Fig. 1b). Overall, 100% of the HiFi and 99.01% of the ultralong ONT reads could be aligned to *T2T-goat1.0*, and the unaligned long reads were found with highly repetitive sequences of two nucleotides [e.g., (AT)_n_], or with failed sequencing due to overloaded errors. In total, 99.92% of the short reads could be aligned to *T2T-goat1.0*, and the remaining unaligned short reads were confirmed to be bacterial contaminants or sequencing errors. The alignments of the Bionano optical maps also supported the accurate assembly of *T2T-goat1.0* (Supplementary Fig. 4).

**Table 1.**
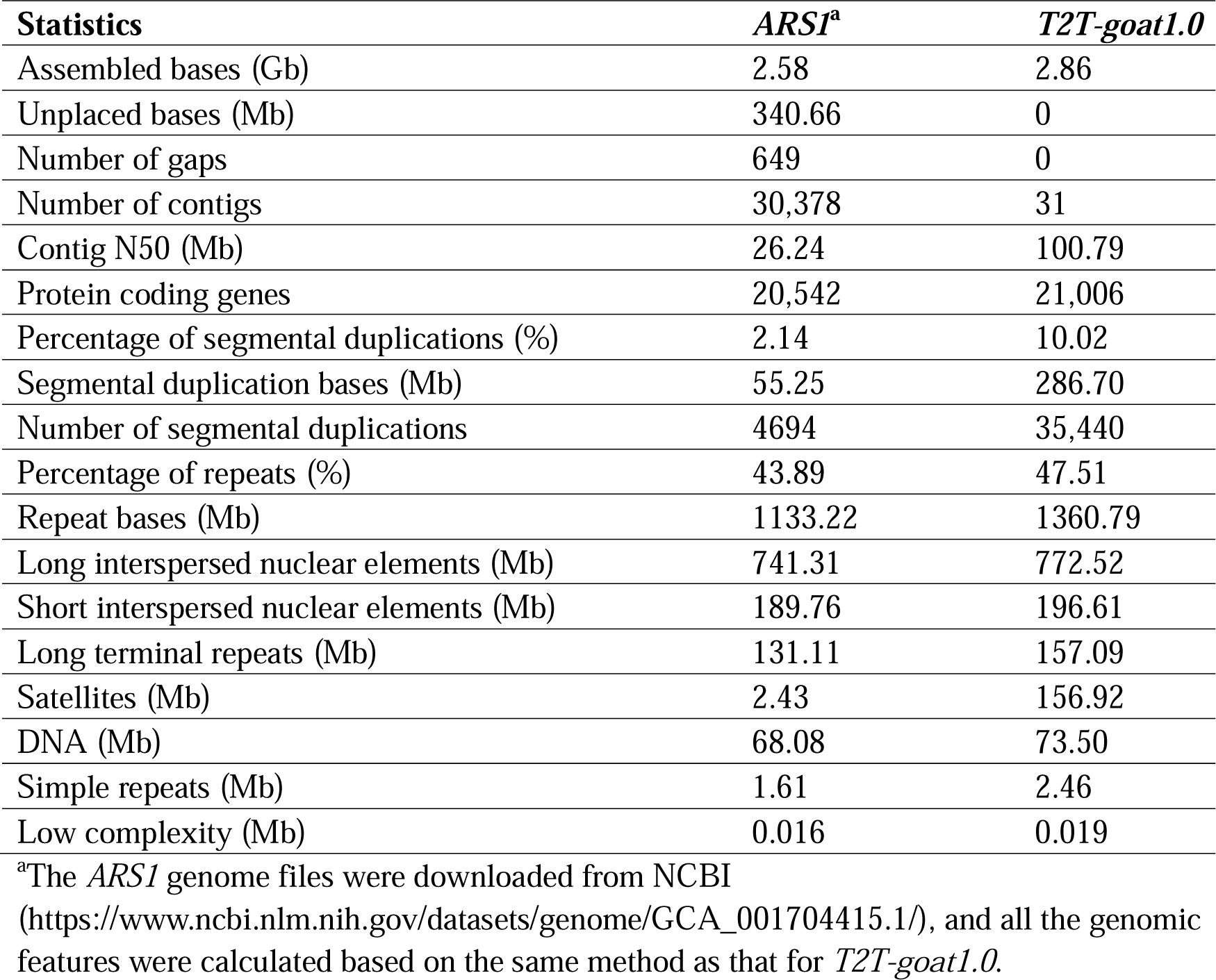
Summay of *T2T-goat1.0* and *ARS1* goat assemblies.

*T2T-goat1.0* represented the longest gap-free chromosome-level genome in all the available goat genome assemblies (Fig. 1c), only two of which included incomplete Y chromosomes. The recent goat assemblies (e.g., GCF_001704415.1, GCA_015443085.1 and GCA_026652205.1) harbored 136 to 649 gaps, 231.45 to 287.20 Mb less in length than the *T2T-goat1.0* assembled here.

### Assembly improvement of *T2T-goat1.0*

High linear synteny was found between *T2T-goat1.0* and the NCBI goat genome reference *ARS1* (GCF_001704415.1), indicating the high quality and reliability of *T2T-goat1.0* (Fig. 2a). The assembly *T2T-goat1.0* closed all the 649 gaps in *ARS1* and resolved the telomeric regions. Many previously unresolved regions (PURs) were constructed by comparing the chromosomes of *T2T-goat1.0* and *ARS1*, for a total size of 288.5 Mb (Fig. 2b). More than 30 Mb of PURs were found on the X chromosome (Fig. 2b). These PURs consisted mostly of centromeric satellites and segmental duplications (SDs) which accounted for 81.92% (236.33 Mb) (Fig. 2c). Additionally, we detected an overall size of 157.38 Mb for the newly assembled regions (NARs) that are not included in *ARS1* and found that most NARs were in centromeric regions. In contrast to the average QV values of 40.93 and 32.83 for chromosomes and all the sequences (including chromosomes and unplaced contigs) in *ARS1* respectively, the QV estimate was greatly enhanced for *T2T-goat1.0*, for which the average QV was 54.19 and the maximum QV was 59.31 for CHI10. Throughout the *T2T-goat1.0* genome, the minimum unique *k*-mer (MUK) was defined as a *k*-mer present only once in the genome with a sliding window of 100 kb, and we observed a substantial increase in the number of unique sequences in *T2T-goat1.0* compared to that in *ARS1* (Supplementary Fig. 5). Longer MUKs in 100-kb windows across the chromosomes were observed in the centromeric regions (Fig. 1d and Fig. 1e).

**Fig. 2.**
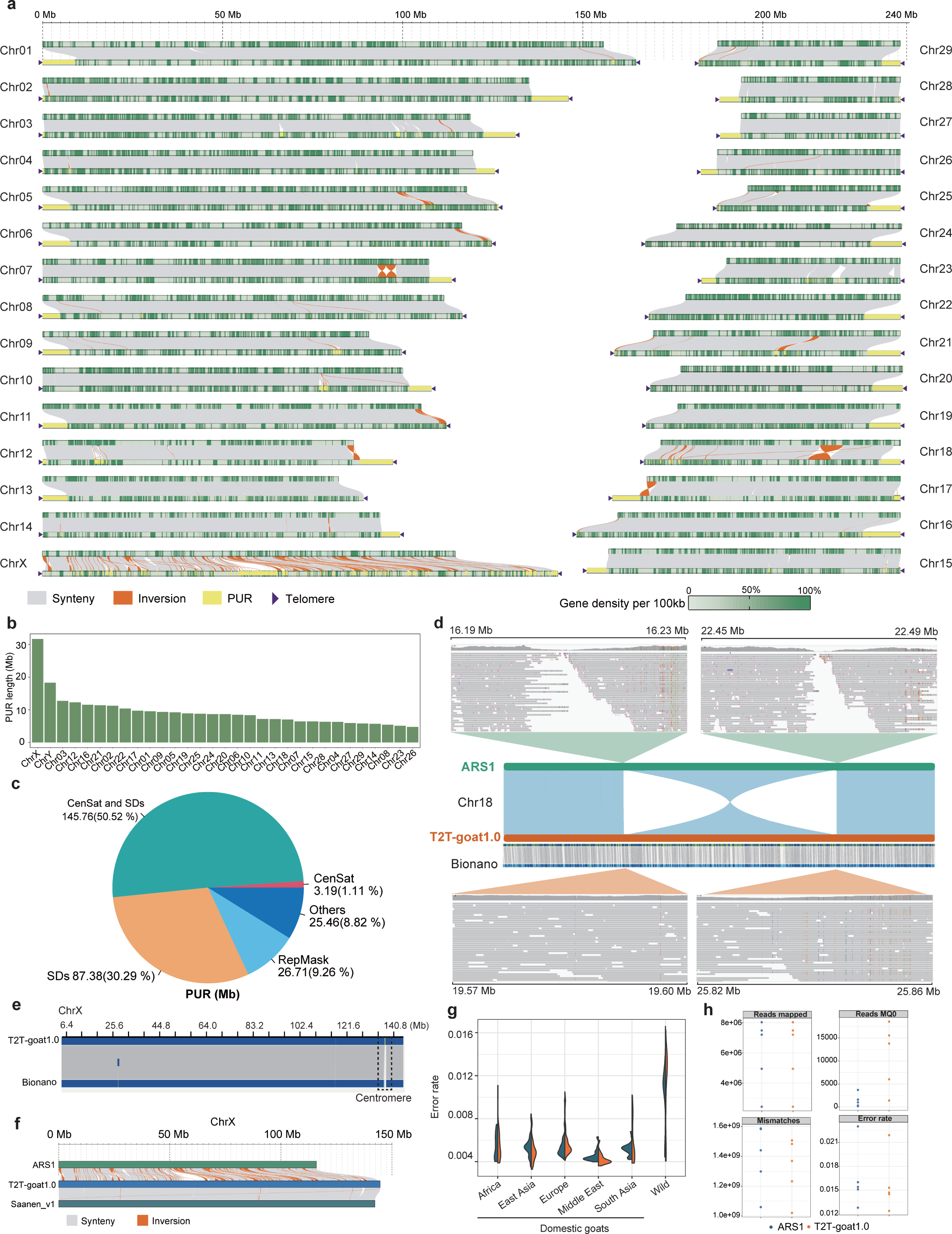
Comparisons of the *ARS1* and *T2T-goat1.0* goat assemblies. **a,** Syntenic and nonsyntenic regions between *ARS1* (top) without telomeres and *T2T-goat1.0* (bottom) with telomeres that are indicated by dark purple triangles. The collinearity between two genome assemblies is shown as gray lines, and the inversions are shown as orange lines. The yellow bars represent the previously unresolved regions (PURs) in *T2T-goat1.0*. The gene density in 100-kb windows is shown as dark green bars. **b,** The total bases of PURs for each chromosome in *T2T-goat1.0* compared to that in *ARS1*. **c,** The proportions of various repetitive elements in PURs. CenSat, satellite sequences in the centromeric region; SDs, segmental duplications; RepMask, RepeatMasker. **d,** An structural error on chromosome 18 of *ARS1*. The two junction sites of an inversion on chromosome 18 of *ARS1* could not be covered by the HiFi reads, and the clipped reads are visualized with IGV, while the corresponding sites in *T2T-goat1.0* were assembled correctly with even long read coverage and Bionano optical maps. **e,** Bionano optical maps of the X chromosome in *T2T-goat1.0*. **f,** Collinearity of the X chromosome among *ARS1*, *Saanen_v1.0*, and *T2T-goat1.0*. Synteny and inversion are shown in gray and orange respectively. **g,** Error rate for alignments of short reads in 5 geographic domestic goat populations and wild goats using *T2T-goat1.0* and *ARS1* as references. **h,** Alignment indices of long reads from five newly sequenced goats using *T2T-goat1.0* and *ARS1* as references.

Additionally, *T2T-goat1.0* corrected abundant structural errors in *ARS1*, including large inversions on CHI07 and CHI18 (Fig. 2a). We identified an inversion on the right arm of CHI18 in *ARS1* compared to *T2T-goat1.0*, and examined the two junction sites via read alignment. We found that the two junction sites were not covered by any long reads sequenced for the *ARS1* genome assembly, but the failed alignment of the reads at the junction sites was rescued by the corresponding region of *T2T-goat1.0* (Fig. 2d). The assembly quality and reliability of this region were supported by the Bionano evidence (Fig. 2d). Another example of the two neighbor inversions on CHI07 in *ARS1* was also confirmed as structural errors (Supplementary Fig. 6). The accurate assembly of the X chromosome was further supported by read alignments (Fig. 1b) and Bionano maps (Fig. 2e). *T2T-goat1.0* filled numerous gaps and corrected multiple structural errors in the highly fragmented sequences on the X chromosome of *ARS1* (Fig. 2f). These structural errors on the X chromosome in *ARS1* were not found in *Saanen_v1* (CM027079.1), and the evaluation of collinearity between *T2T-goat1.0* and *Saanen_v1* demonstrated only 8 inversions (Fig. 2f). In addition to the Y chromosome, our *T2T-goat1.0* had 4.62 Mb and 31.68 Mb of PURs added to the X chromosomes of the *Saanen_v1* and *ARS1* assemblies, respectively.

Furthermore, we investigated the reference bias of read alignment for population genetics analysis. Short reads from NGS were retrieved from worldwide domestic and wild goats (Supplementary Table 3). Compared to *ARS1*, *T2T-goat1.0* enabled confident mapping of short reads in terms of more reads mapped and a lower mismatch and error rate (Fig. 2g). *T2T-goat1.0* improved the number of mapped reads significantly, and >30.91% of the goat samples had >1% more short reads aligned (19.04% with >5% more reads and 1.98% with >10% more reads) than to *ARS1*. We observed an increased number of MQ0 reads in *T2T-goat1.0*, which could be attributed to the fact that most of the newly added regions are composed of repetitive sequences, and short reads are not advantageous in aligning to repetitive sequence positions due to their short lengths. Short reads from the Middle East and East Asia populations showed significantly lower numbers of mismatches and fewer errors, benefiting from their relatively close genetic relationships with the Inner Mongolia cashmere goat^2^, the goat population with its T2T genome assembled here (Fig. 2g and Supplementary Fig. 7). Moreover, we sequenced the genomes of five goats from five representative populations to produce long reads (Supplementary Table 1). Like for short reads, more aligned reads and lower mismatch and error rates were observed in the alignment of the long reads against *T2T-goat1.0* than against *ARS1* (Fig. 2h and Supplementary Table 4).

### Genome annotation

Annotation of repeat content based on a combination of prediction and evidence-based tools revealed 47.51% repetitive elements, including 26.97% long interspersed retrotransposable elements (LINEs), 6.87% short interspersed retrotransposable elements (SINEs), 5.48% long terminal repeats (LTRs) and 5.48% satellites (Fig. 1d, Fig. 1e, Supplementary Fig. 8, and Table 1). Enrichment of satellites was present in the centromeric region, with rare satellite repeats found on the Y chromosome, although LINEs and SINEs were widely distributed on each chromosome (Supplementary Fig. 8 and Supplementary Fig. 9). The repetitive sequences of centromeric satellites, SDs, and repeats identified by RepeatMasker software dominated the PURs, with a total length of 263.04 Mb (91.18%) (Fig. 2c). Overall, the repetitive sequences in PURs made up 12.91% of the entire genome’s repetitive sequences and occupied 94.92% of all the satellite sequences. LTRs were found to be distributed across all chromosomes, preferably with peaks around centromeric regions enriched with satellites (Supplementary Fig. 9).

A combined strategy of *de novo*, transcriptome-based and homolog-based prediction was used to perform gene annotation. A total of 21,006 high-confidence protein-coding genes were obtained (Table 1) with the average gene length of 46.97 kb, compared to 44.53 kb in *ARS1*, and 97.90% of the Benchmarking Universal Single-Copy Orthologs (BUSCO)^23^ were annotated. Among these genes, 1126 were newly anchored genes in PURs, and 446 were newly assembled genes (NAGs) in NARs that were not included in *ARS1*. RNA-seq of 14 goat tissues (Supplementary Table 1) supported the expression of these novel genes, and NAGs were diversely expressed in tissues of the liver, spleen, rumen, etc. (Supplementary Fig. 10a). We further annotated the functions of NAGs against the available protein databases, such as NCBI’s nonredundant proteins (NR) and Gene Ontology (GO) databases. The NAGs in the NARs were significantly (*P*_adj_ < 0.05) enriched in the pathways (Supplementary Fig. 10b) related to GTP binding, protein deubiquitination, cysteine-type deubiquitinase activity, etc. Additionally, no genes were annotated within the centromeric regions of autosomes or the X chromosome. A tangle of the assembly string graph located far from the centromeric end was found to be associated with 45S rDNA repetitive sequences, comprising of 18S, 5.8S, and 28S on five chromosomes, CHI2, CHI3, CHI4, CHI5, and CHI28, which was confirmed by fluorescence in situ hybridization (FISH) signals (Supplementary Fig. 3). The 5S rDNA was only found on the right end of CHI28, opposite to the 45S rDNA end, as shown in both the assembly and FISH. The results revealed a pattern of telomere (T) - rDNA (R) – centromere (C) – telomere (T) (TR-CT), with centromere and rDNA located on either of the chromosomal ends.

We calculated the sequence length (>1 kb) and alignment identity (>90% identity) to determine the SDs of *T2T-goat1.0*. We identified a total of 286.70 Mb nonredundant SDs in *T2T-goat1.0*, while only 55.25 Mb was found on the chromosomes of *ARS1*. *T2T-goat1.0*showed a remarkable increase in the number and length (additional sequences of 212.18 Mb) of interchromosomal SD pairs compared to those of *ARS1*. Overall, 275 genes in 118 families overlapped the regions with SDs, indicating a critical contribution of SDs to the expanded gene families. Furthermore, we observed significantly greater gene copy numbers in *T2T-goat1.0* than in *ARS1* and the goat assembly *ASM2665220v1* (GCA_026652205.1), which could be attributed to the successful assembly of repetitive sequences and genome annotation improvements in *T2T-goat1.0* (Supplementary Table 6). For example, in the gene family associated with scavenger receptor activity (OG0000004), 15 gene copies were annotated on CHI05 in *T2T-goat1.0* instead of only 8 in *ARS1* (Supplementary Fig. 11).

### Genome structure of centromeres and telomeres

We focused on deciphering the centromeric structure and repetitive structure in the unique acrocentromeric chromosomes (autosomes and X chromosome) and metacentric Y chromosome. First, we determined the centromeric regions based on the enriched regions of the mapped ChIP-seq reads with the histone antibody Phospho-CENP-A (Ser7) and the enriched methylated cytosine sites based on the HiFi and ONT data (Fig. 3a). Furthermore, the centromeric regions on the acrocentric autosomes were confirmed by a combination of ChIP-seq peaks, DNA hypermethylation, Hi-C interaction heatmaps, and enriched satellite DNAs (Supplementary Fig. 12a and Supplementary Fig. 12b). The lengths of the centromeric regions ranged from 0.60 Mb to 8.51 Mb (Fig. 3b), with a short centromere (0.60 Mb) on the X chromosome. Repeat annotation of centromeric regions revealed the dominance of satellite DNAs, and to identify high order repeats (HORs), 21 centromeric satellite sequences were identified using the SRF software^24^. Pairwise comparisons classified the 22 repetitive sequences into three categories of centromeric satellite units, i.e., SatI (816 bp), SatII (702 bp), and SatIII (22 bp).

**Fig. 3.**
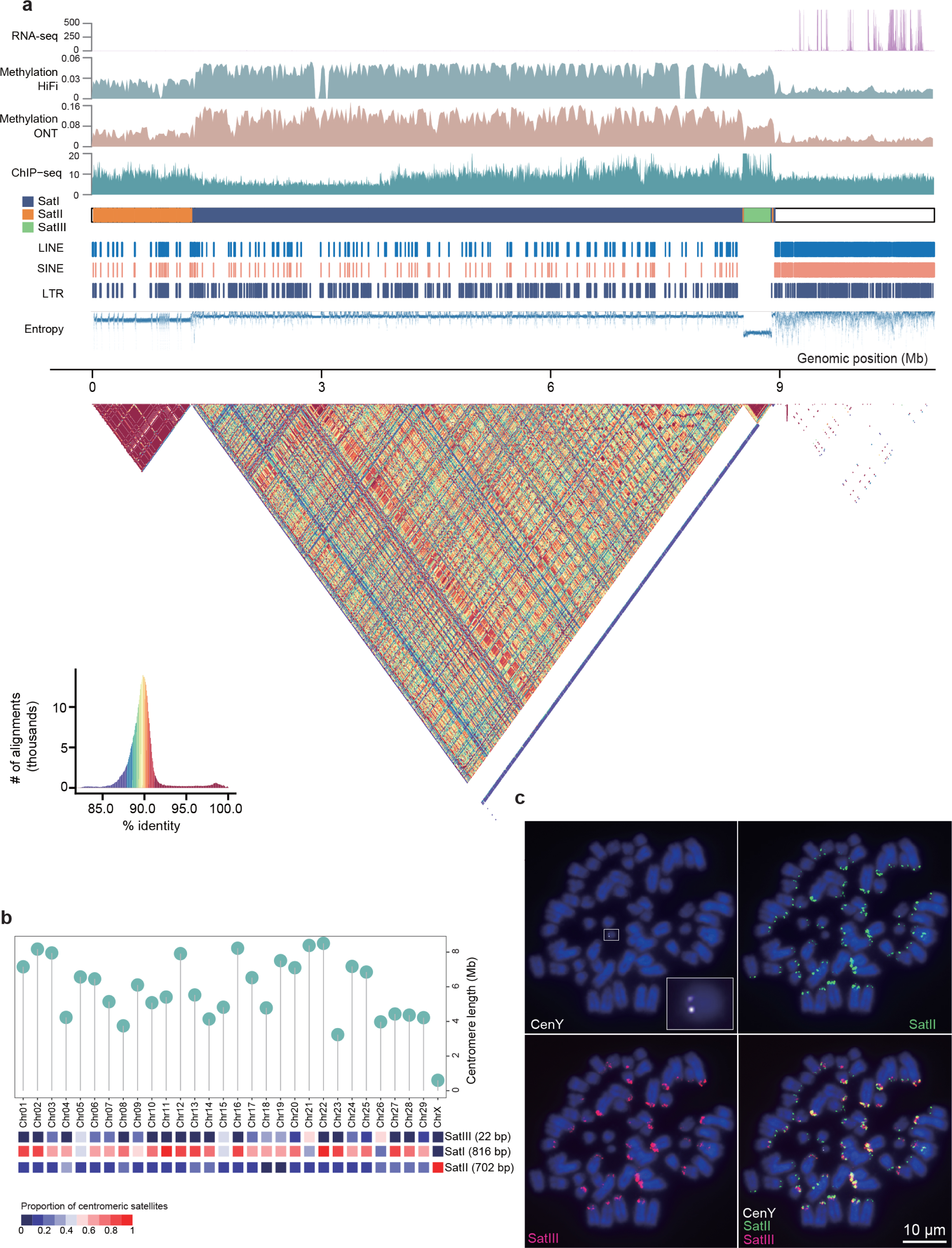
Genomic structure of centromeric regions. **a,** Schematic representation showing the sequence compositions in the centromeric region of chromosome 1. From top to bottom: RNA-seq, expression (read counts in 1-kb windows) of genes in the pericentromeric region; Methylation by HiFi and ONT in 20-kb windows; ChIP-seq for CENP-A protein enrichment based on Phospho-CENP-A (Ser7) antibodies in 5-kb windows; Satellites of SatI, SatII and SatIII; LINE; SINE; LTR; and Entropy that was calculated for sequence complexity with low values for centromeric regions. The pairwise 10-kb sequence identity (%) heatmap in centromeric region is shown below. **b,** Length distribution of centromeric regions and proportions of three centromeric satellite units of all the autosomes and the X chromosome. **c,** FISH results for probes of CenY (white), SatII (green) and SatIII (red). CenY is shown uniquely on the Y chromosome.

Satellite compositional bias was detected across the chromosomes (Fig. 3b). The most abundant centromeric satellite repeat, SatI, was found in the centromeric regions of all autosomes and the X chromosome (Fig. 3b) and had an accumulative length of 118.19 Mb, covering approximately 67.81% of their centromeric regions. SatI was previously identified to be a constant length of 816 bp (X57335.1) and could be mapped well to the centromeric regions of *T2T-goat1.0*. SatII accounted for 15.21% of the total satellite sequences in *T2T-goat1.0*, with a total length of 26.50 Mb. Notably, the genomic locations of SatII were always associated with ChIP-seq peaks in the centromeric regions. For example, SatII was found to cooccur with ChIP-seq peaks on CHI22, CHI28, and the X chromosome. However, a specific link between SatII and ChIP-seq was not implied because ChIP-seq peaks could also be found in centromeric regions containing SatI, for example, on CHI20 and CHI21. We observed a high consistency on the occurrence of SatII across all the chromosomes except the Y chromosome in both FISH and the genome assembly (Fig. 3c). In contrast to the most abundant SatI and SatII, SatIII was not typically present in the core centromeric region but was present within the centromeric near-end regions, suggesting the evolutionary history of centromeric regions. SatIII contains two major variants differing in two nucleotides, which showed the intensified FISH signals of probe binding on different chromosomes (Fig. 3c and Supplementary Fig. 12c). We confirmed the conservation of centromeric satellite sequences between sheep and goats since the ovine centromeric SatI (KM272303.1, 816 bp^25^) and SatII (U24092.1, 440 bp^26^) sequences could be mapped well to the *T2T-goat1.0* due to their high similarity.

To further understand the centromeric evolution on each chromosome, we generated pairwise sequence identity heatmaps within and around centromeric regions, which included distinct classes of HORs related to SatI, SatII, and SatIII. Compared to those of SatI, higher identity values for SatII and SatIII were observed, indicating closer similarities and fewer variations in the repeat sequences possibly associated with their recent origins (Supplementary Fig. 12a). This observation suggested centromeric reorganization of HORs, which was in accordance with the results of sequence complexity analysis (Entropy, Fig. 3a). For example, we found three major evolutionary layers corresponding to SatI, SatII and SatIII in the sequence identity heatmap of centromeric region of CHI1, and minor footprints of SatI and SatII around the SatIII layer (Fig. 3a).

We noted that the methylated CpG sites identified by HiFi and ONT reads showed a dip inside the centromeres, where the ChIP-seq signals were enriched, for example, SatII HORs on CHI01 (Fig. 3a) and CHI06 (Supplementary Fig. 12a). The hypomethylation of goat centromeric regions is similar to that in humans^10,27^, and the overlap between hypomethylated regions and ChIP-seq signals suggested interactions between methylation and histone activities. Hypomethylation in centromeric regions promotes CNEP-A deposition and binding of functional kinetochores^27,28^. Additionally, an invasion of other repeats, including LINE and SINE, was detected in the centromeric regions; notably, a LTR (4852 bp) was distributed across nearly all the centromeres, with an accumulative length of 1.36 Mb.

By employing a sequence query based on the telomeric repeat (TTAGGG at the 5’ end and CCCTAA at the 3’ end), we uncovered 62 telomeres in the assembly spanning 0.71 Mb. This led to the creation of 31 telomere-to-telomere chromosomes. The length of the identified telomeres ranged from 2,993 to 26,638 nucleotides across all chromosomes, with the longest telomere located at the right end of CHI3 (Supplementary Fig. 13).

### Y-chromosome assembly and structure

We used paternal-specific ONT ultralong reads (2.26 Gb, >100× coverage) to assemble the Y chromosome. The complete Y chromosome (*T2T-CHIY1.0*), with a length of 20.9 Mb, presented an overall QV of up to 49.10 after polishing, and the mapped HiFi and ONT reads showed evenly distributed coverage (Fig. 1b). The two available Y chromosome assemblies of *Saanen_v1*^6^ (CM027080.1) and *ARS1*-based scaffolded contigs^29^ (GCA_018357025.1) are only 9.6 Mb and 11.5 Mb in length, respectively, and were used to evaluate *T2T-CHIY1.0* quality. According to the collinearity plots, the syntenic regions of ∼7 Mb between the left arms of chromosomes X and Y were not assembled in these two incomplete Y chromosome assemblies (Supplementary Fig. 14). The SD-enriched region on the *T2T-CHIY1.0* coordinates of 13∼19 Mb (Fig. 4a) resulted in poor collinearity, referring to low-identity organization with multiple contigs in the Y chromosome of *ARS1* (Fig. 4b) and abundant inversions in the Y chromosome of *Saanen_v1* (Fig. 4c). We found 24 gaps in this region with poor-collinearity on Y chromosome of *Saanen_v1*, which could be ascribed to the presence of gene arrays as shown in *T2T-CHIY1.0*.

**Fig. 4.**
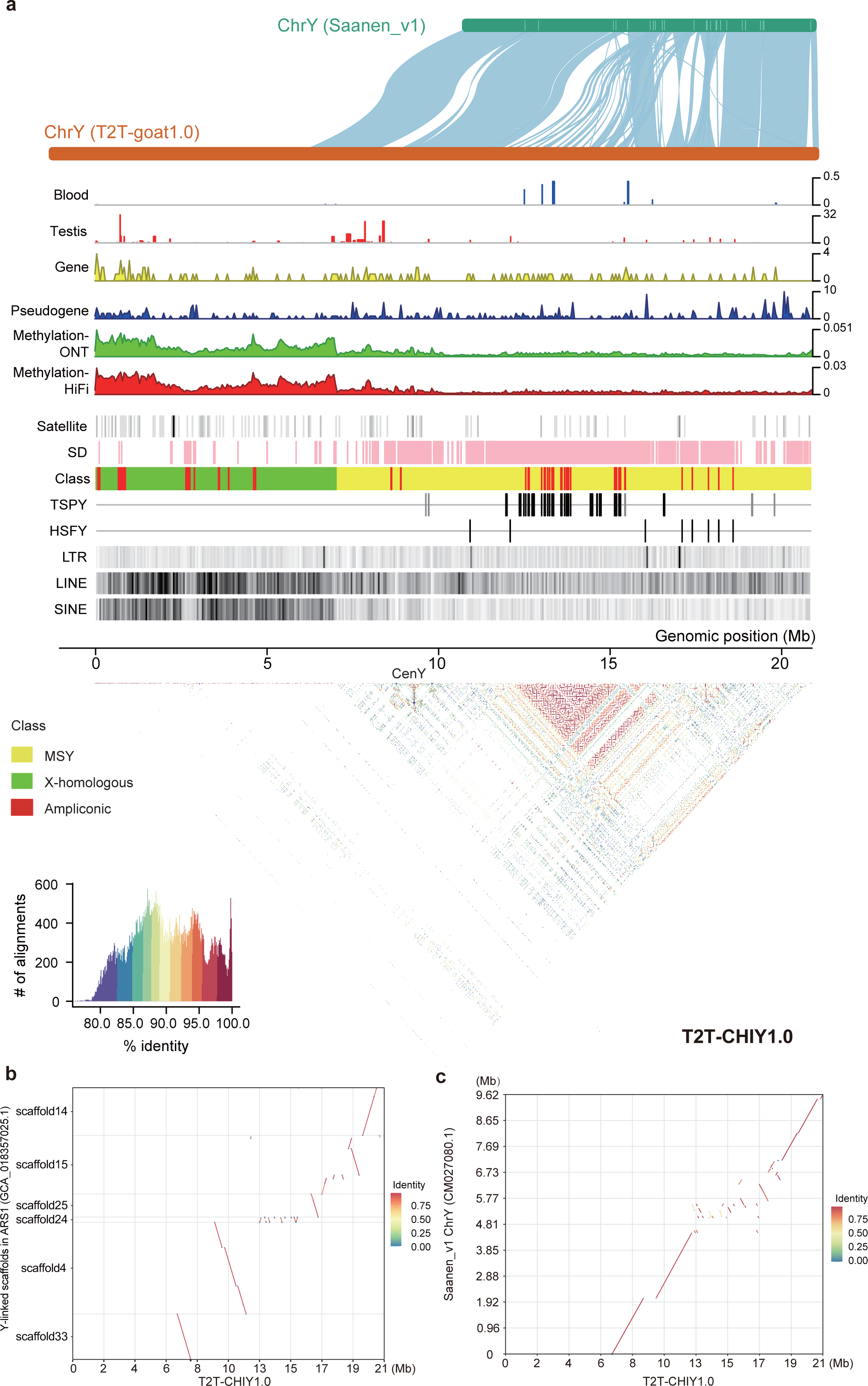
Genomic structure of the Y chromosome. **a,** Genomic structure and features of the Y chromosome in *T2T-CHIY1.0*. From top to bottom: collinearity of Y chromosomes between *Saanen_v1.0* and *T2T-goat1.0*; Gene expression in blood and testis tissues; Protein-coding gene density; Pseudogene density; Methylation (5mC) levels estimated with ONT and HiFi reads; Satellites; Segmental duplications (SDs); Class, X-chromosome-homologous (green), male-specific Y (MSY, yellow) and ampliconic (red) regions are included in *T2T-CHIY1.0*; *TSPY* genes highlighted for Clade I in black and Clade II in gray; *HSFY* genes; LTR density; LINE density; and SINE density. The density of methylation, Satellite, LINE, SINE, and LTR is shown in 50-kb windows. The pairwise 10-kb sequence identity (%) heatmap across *T2T-CHIY1.0* is shown below. **b,** Collinearity of the Y chromosomes of *ARS1* and *T2T-CHIY1.0*. **c,** Collinearity of the Y chromosomes of *Saanen_v1.0* and *T2T-CHIY1.0*.

*T2T-CHIY1.0* unlocked the PURs of 7.68 Mb compared to the *Saanen_v1* Y chromosome, which mostly fell into the homologous region between chromosomes X and Y (Fig 4a). By annotating the repetitive elements on the Y chromosome, we found that the repetitive content reached 53.79%, while the repetitive content on autosomes was only 45%. The most abundant repetitive sequences were LINEs (7.42 Mb in length), which accounted for 35.5% of the entire repetitive sequence, followed by the LTR elements (1.82 Mb, 8.70%) and SINE elements (1.42 Mb, 6.79%). Furthermore, LINE and SINE elements are more densely distributed in the X-chromosome-homologous region on the left arm of the Y chromosome.

We annotated 129 protein-coding genes and 185 pseudogenes that were diversely distributed across the whole Y chromosome, of which 54 genes were newly found in the PURs of *T2T-CHIY1.0* (Fig 4a). The annotation revealed two or more copies of 74 genes in the ampliconic regions on CHIY (Supplementary Table 7), including 27 gene copies of testis-specific protein Y-encoded (*TSPY*) and 8 copies of heat shock transcription factor Y-linked (*HSFY*) (Fig. 4a). Nevertheless, only one copy of *TSPY* and two copies of *HSFY* were detected in *ARS1*. The *TSPY* genes on the Y chromosome were expressed only in the testis and were clustered in the region with enriched SDs, which was confirmed and supported by the sequence identity heatmap (Fig. 4a). We constructed a phylogenetic tree of *TSPY* genes using human *TSPY* genes as the outgroup (Supplementary Fig. 15). The results revealed that the goat *TSPY* family could be divided into two clades. Clade I, with 21 *TSPY* copies, exhibited a close sequence distance as a cluster in a continuous region of ∼3.1 Mb. The six gene copies in Clade II surrounded those in Clade I on *T2T-CHIY1.0*, which suggested the two-step patterns of *TSPY* amplification. Similar to *TSPY*, the repeat array of *HSFY* was located in the SD-enriched region (Fig. 4a). Furthermore, we annotated 32 single-copy genes, including Y-chromosome-specific genes such as *SRY*, *UTY*, *DDX3Y*, and *USP9Y*.

Based on the HiFi data, hypermethylation in blood was identified on the left arm of *T2T-CHIY1.0*. This suggested that the genes in this region were preferentially not expressed in blood due to transcriptional inhibition caused by hypermethylation of the genes’ bodies and promoters. As SatI, SatII and SatIII were not found on the Y chromosome, we extended our search for repeat sequences on the sheep Y chromosome. A new repeat sequence of 1474 bp was inferred as the putative centromere-specific repeat unit, CenY, since it was highly homologous to CenY (2516 bp) in the centromeric region on the Y chromosome of *T2T-sheep1.0* (T2T genome assembly for a ram of the Hu sheep breed). CenY was enriched in the middle of the Y chromosome, i.e., at the border of the hypermethylated region, which was indicated by a single sharp signal in the sequence identity heatmap, and CenY was not present in any of the autosomes or the X chromosome as shown in sequence search of *T2T-goat1.*0 and FISH results (Fig. 3d).

We tested the performance of *T2T-CHIY1.0* as a reference to call variants for a population of 102 bucks (Supplementary Table 3), which yielded a mapping rate as high as 99.95%. The average alignment rate using *T2T-goat1.0* as a reference reached 98.07% in the buck population, whereas it was 97.96% using *ARS1*.

### Structural variants based on long reads

To assess the advance of *T2T-goat1.0* as a reference for long-read alignment and calling of large-scale structural variations (SVs), we used the PacBio Sequel II platform to sequence the genomes of five female goats of five representative breeds with diverse origins (Supplementary Table 1 and Supplementary Fig. 16a). After quality control, we aligned the long reads from these five individuals to both *T2T-goat1.0* and *ARS1* and called SVs for comparison between the two assemblies. On average, we identified a total of 63,417 SVs in the PacBio long reads of the five individuals using *T2T-goat1.0* as a reference. The number of SVs increased approximately 1.25-8.89% compared with that of *ARS1* (Fig. 5a). The increased in SVs was attributed mostly to deletions and insertions rather than the inversions and translocations. Compared with that of *ARS1*, the use of *T2T-goat1.0* as a reference genome led to the identification of slightly more deletions than insertions (Fig. 5b and Fig. 5c). The lengths of the deletions and insertions exhibited a distribution peak of 1-2 kb (Supplementary Fig. 16b and Supplementary Fig. 16c), which is longer than that of 200-300 bp in the human *T2T-CHM13* genome^18^. The improvement in the detection of deletions and insertions could be attributed to the accurate assembly of repetitive sequences in *T2T-goat1.0*. We discovered a substantial increase in deletions and insertions within the satellite sequences, with more than 9,000 by *T2T-goat1.0* (Fig. 5b) versus 359 by *ARS1* (Fig. 5c); however, no significant increase in the other repetitive sequences, such as LINEs, SINEs, or LTRs, was observed (Supplementary Fig. 16d and Supplementary Fig. 16e). The improvement in SV detection resided not only in the quantity of deletions and insertions on satellites but also in the length frequency shift, with peak lengths of 751-1000 bp in *T2T-goat1.0*, compared to 50-100 bp in *ARS1*.

**Fig. 5.**
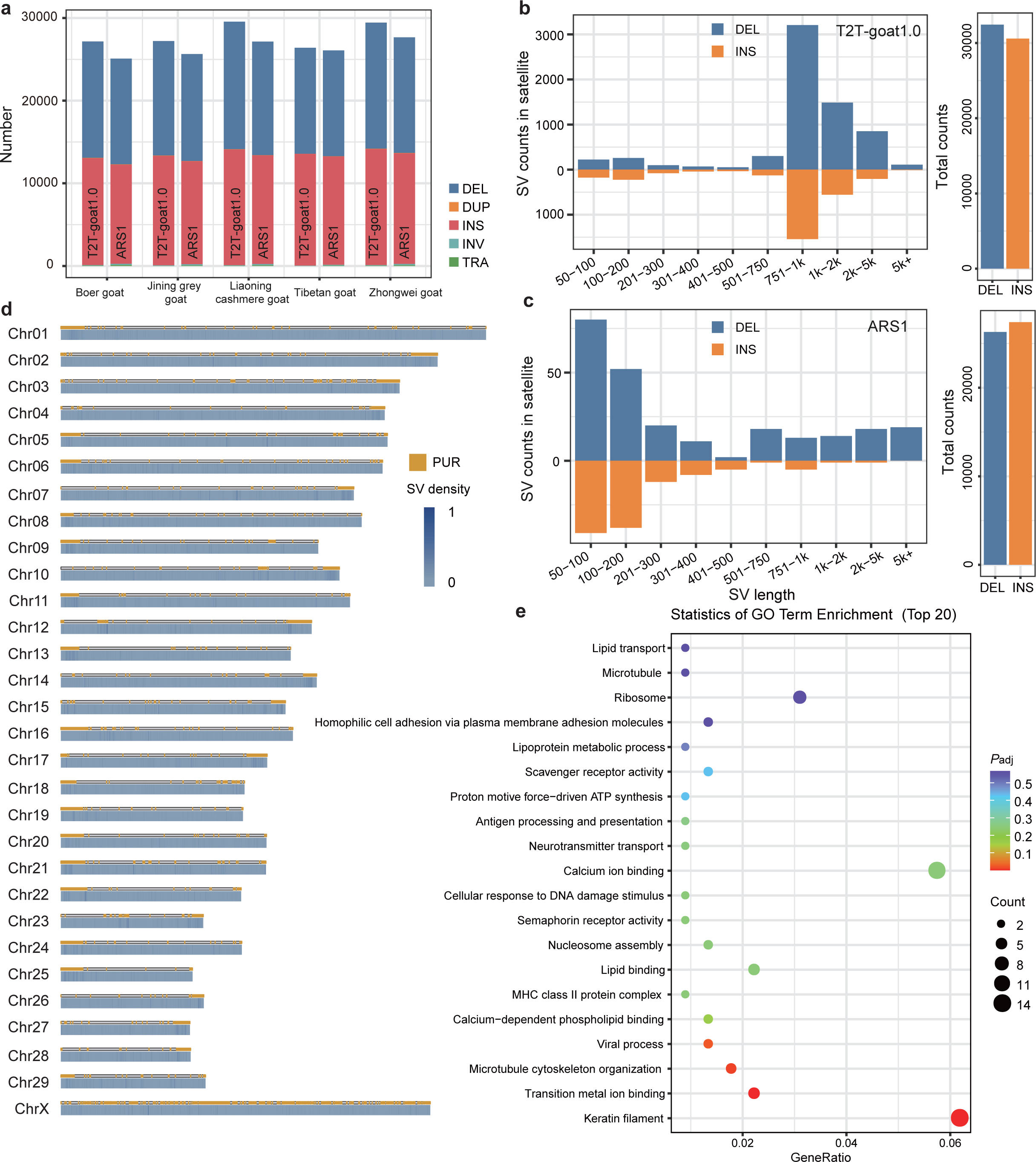
Improvements in structural variants based on long reads. **a,** *T2T-goat1.0* and *ARS1* as references are compared for performances of calling SVs based on long reads of five goats (BEG, JNGG, LNCG, TG, and ZWG). DEL, deletion; DUP, duplication; INS, insertion; INV, inversion; TRA, translocation. Counts of deletions and insertions were summarized in the satellite sequences in a comparison of *T2T-goat1.0* (**b**) and *ARS1* (**c**) as references. The total numbers of deletions and insertions are shown on the right. **d,** Density of SVs called from PacBio reads with *T2T-goat1.0* as a reference, in 10-kb windows. PURs are highlighted in yellow. **e,** GO enrichment of genes whose exons overlapped with the SVs called with *T2T-goat1.0* as a reference.

The accurate assembly and completeness of the PURs in *T2T-goat1.0* enabled us to resolve complex genomic regions and enhance SV calling (Fig. 5d). Among the total SVs, up to 4711 SVs (7.41%) were identified to cluster densely in the PURs, particularly in the centromeric regions, consisting of 3021 deletions and 1690 insertions. We found that 18,804 SVs fell into the gene bodies, including 507 SVs overlapping exons. GO analysis revealed that these 507 genes were enriched in the keratin filament pathway (Fig. 5e), suggesting that these SVs are crucial in fiber formation and development in cashmere goats^30,31^. We further examined the 57 SVs that uniquely occurred in *T2T-goat1.0* but were not in the assemblies of the other five goats, and found that *FGFBP1*^32,33^, *KAP9-2*^34^ and *COL1A1*^35^ are known to be associated with hair and cashmere development (Supplementary Table 8). A total of 5224∼6584 SVs were uniquely identified in the five female goats (Supplementary Fig. 16f), and covered exons in 40∼49 genes (Supplementary Table 9), which are frequently involved in olfactory receptors, interferon, immune system, and cashmere development. For example, mucin (MUC) family genes, including *MUC2*, *MUC5B* and *MUC6*, on CHI29 experienced the independent deletions in the four goats except for Tibetan goat. Two insertions in *TNNI1* gene were found in Tibetan goat, and *TNNI1* is involved in the calcium-sensitive regulation of skeletal muscle contraction, and associated with oxygen level and aerobic exercise^36^.

### New variants based on short reads

To investigate the impact of *T2T-goat1.0* on short-read variant calling, we collected NGS data from 516 caprine samples (∼15x on average), which consisted of 68 wild and 448 domesticated goats from five major geographic regions worldwide (Supplementary Table 3 and Fig. 6a). Clean short reads of the genomes were aligned to both *T2T-goat1.0* and *ARS1* for single nucleotide polymorphism (SNP) calling. After population-wide SNP calling and filtering for quality control, we obtained a total of 25,397,794 high-quality SNPs across the samples against *T2T-goat1.0*, whereas 24,238,138 SNPs were yielded when mapping to *ARS1*. *T2T-goat1.0* added various high-quality SNPs compared to those of *ARS1*, ranging from 6,405 to 67,614 on each chromosome. For example, the number of SNPs on the X chromosome increased from 861,416 to 977,085 (Supplementary Fig. 17a), which could have resulted from the base-level errors in *ARS1* (Fig. 2f). Subsequently, we identified a total of 545,026 SNPs within the PURs (Supplementary Fig. 17b), which accounted for 2.1% of the total SNPs called using *T2T-goat1.0* as a reference. We observed the most SNPs in wild goats, and compared to that with *ARS1* as a reference, we identified an increase of total SNPs in all populations, including homozygous and heterozygous SNPs, except for East Asia (Supplementary Fig. 17c, Supplementary Fig. 17d and Supplementary Fig. 17e).

**Fig. 6.**
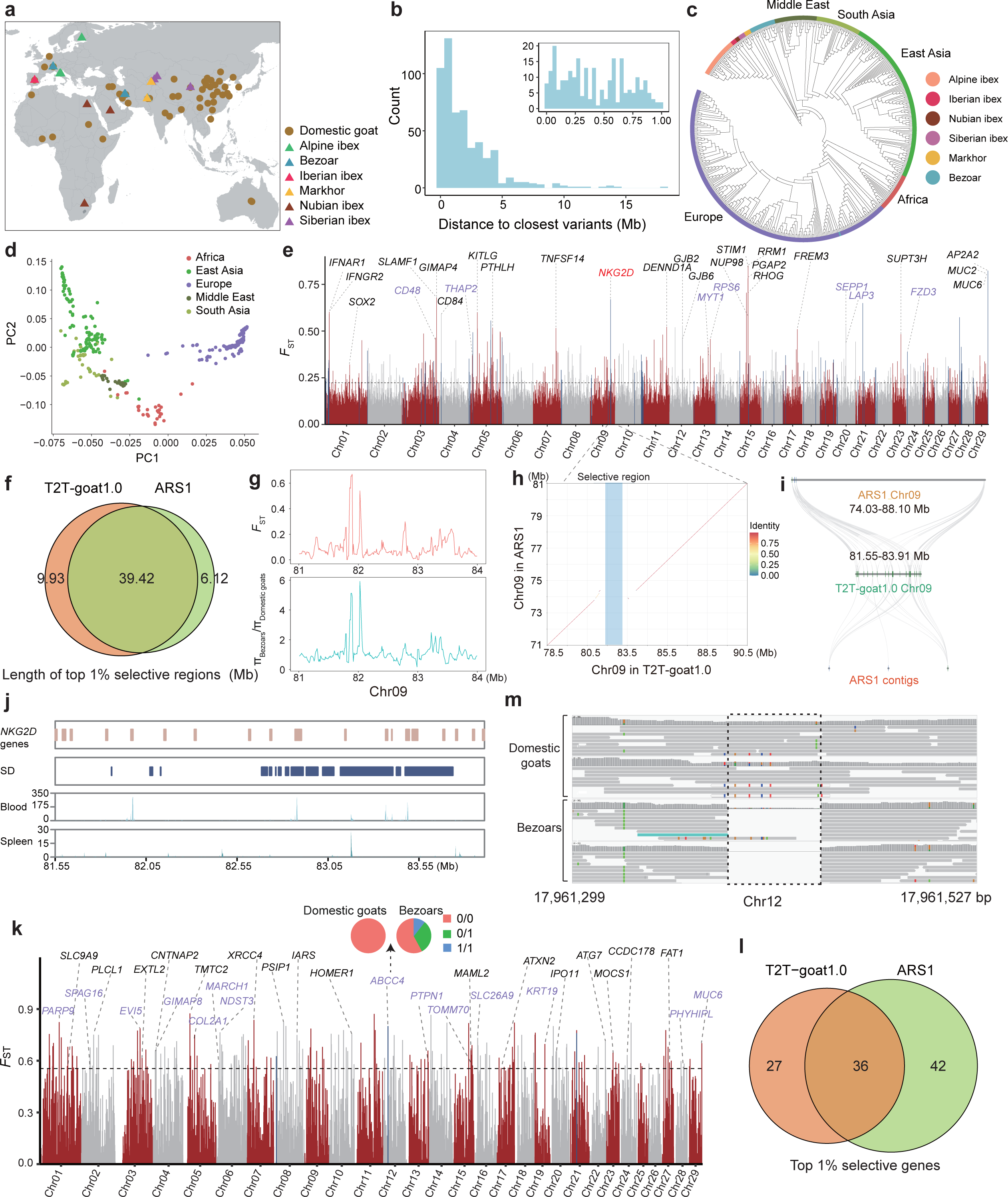
Population structure and selection signatures for domestication. **a,** Geographic distributions of domestic goats and wild goats. **b,** Distance of the closest variants to the goat QTLs, with the distance of <1 Mb shown in the right top corner. The neighbor-joining tree of wild and domestic goats (**c**) and PCA of five geographic domestic goat populations (**d**) were generated based on LD-pruned SNPs. **e,** Genome-wide selective signals based on SNPs and *F*_ST_ between bezoars and the domestic goats. The selective signals in the PURs are highlighted in blue bars. The gene symbols are shown in black for the ones identified by both *T2T-goat1.0* and *ARS1*, and the ones only by *T2T-goat1.0* in purple. *NKG2D* in red uniquely identified by *T2T-goat1.0* is selected to be demonstrated as followed. **f,** Venn plot of selective regions based on top 1% *F*_ST_ values between *T2T-goat1.0* and *ARS1* as references. **g,** Selective region with tandem *NKG2D* genes based on the *F*_ST_ and π ratio. **h,** Syntenic plotting of the region with *NKG2D* genes on chromosome 9 between *T2T-goat1.0* and *ARS1.* The selective region highlighted in blue is exactly located inside the region where tandem *NKG2D* genes are not assembled well in *ARS1*. **i,** Collinearity of the region with tandem *NKG2D* genes on chromosome 9 between *T2T-goat1.0* and *ARS1*. The tandem *NKG2D* genes in *T2T-goat1.0* were not placed only on chromosome 9 of *ARS1*, but some of them were scattered on the three unplaced contigs. **j,** Tandem *NKG2D* genes are in accordance with SDs and showed their expression in blood and spleen. **k,** Selective signals based on SVs and *F*_ST_ between bezoars and the domestic goats. The gene symbols are shown in the same colors to that of Fig. 6e. **l,** Venn plot of selective genes that overlapped with selective regions based on top 1% *F*_ST_ values identified by *T2T-goat1.0* and *ARS1*. **m,** A deletion was confirmed within *ABCC4* (IMCG12g00097) of bezoars in IGV.

To determine the potential of novel SNPs associated with the available quantitative trait loci (QTLs), we overlaid the SNPs in PURs with the 558 goat QTLs underlying 26 different base traits and 15 trait variants (https://www.animalgenome.org/cgi-bin/QTLdb/CH/index) from the Animal QTL Database (Animal QTLdb)^37^. The distribution of SNPs in the PURs was <5 Mb from the closest QTLs, and 241 SNPs with a distance of <1 Mb further suggested their potential association with candidate traits (Fig. 6b).

Additionally, we identified a total of 32,419 SVs based on short reads, including 23,466 deletions, 2284 duplications, 1552 inversions, and 5117 translocations, among which 870 new SVs were found in the PURs. Overall, 65.84% of these SVs were in intergenic regions, and the others within or near genic regions included 4.89% in exonic regions, 27.56% in intronic regions, and 1.58% upstream or downstream of genes.

### Population genetics and domestication selection with new variants

SNPs by *T2T-goat1.0* was reliable for elucidating phylogenetic relationships within the genus Capra. A neighbor-joining (NJ) tree was used to divide the entire samples into two major branches, i.e., wild and domestic goats. The domesticated goats were further divided into five clades based on the geographical origin, namely, Europe, Africa, East Asia, South Asia, and the Middle East (Fig. 6c and Supplementary Fig. 18a), and the genetic structure of the global populations was supported by both principal component analysis (PCA, Fig. 6d and Supplementary Fig. 18b) and ADMIXTURE analysis (Supplementary Fig. 18c and Supplementary Fig. 18d). Linkage disequilibrium (LD) analysis indicated that the LD decay rate in wild goats was relatively slow, with the East Asian goat population exhibiting the fastest decay rate compared to the other geographic populations (Supplementary Fig. 18e and Supplementary Fig. 18f). All these results were in accordance with those obtained based on *ARS1*.

With *T2T-goat1.0* as the reference genome, the genetic differentiation statistic *F*_ST_ values in 50-kb windows were calculated based on SNPs as an indicator of domestication signals between the genomes of bezoars (wild goats, *Capra aegagrus*) and the whole domestic goat population (Fig. 6e). Selective sweeps harbored 2631 genomic regions, covering 829 genes (Supplementary Table 10). The selective regions associated with domestication spanned 49.35 Mb in *T2T-goat1.0*, with 39.42 Mb commonly found in *ARS1* (Fig. 6f and Supplementary Fig. 19a). Among them, the signals bore previously reported genes, including *STIM1*, *RRM1*, *MUC6*, *IFNAR1*, *FREM3*, and *CD84*, which underwent strong selection during the domestication process of goats. We narrowed our search to 406 selective signals located in the PURs, spanning 5.93 Mb, and identified 56 genes resulting from our complete assembly of *T2T-goat1.0*. For example, a selective signal of the top 1% on CHI9 (*F*_ST_=0.67) encompassed the tandem genes of *NKG2D*. It serves as an activating receptor and regulator of immune cell responsiveness that can be induced in response to infection by diverse types of pathogens^38^. Remarkably, the signals overlapping with *NKG2D* are supported by the π ratio statistic (Fig. 6g). The 13 SNP loci associated with this selective signal showed significant allelic frequency differences between bezoars and domesticated goat populations (Supplementary Fig. 20a). Accurate assembly around this selective signal resulted in 18 tandem *NKG2D* genes on CHI9 in *T2T-goat1.0*, while the collinearity analysis revealed a loss of large fragments (Fig. 6h) and only 10 *NKG2D* genes located on CHI9 and three other unplaced contigs (Fig. 6i) in *ARS1*. The tandem *NKG2D* genes on CHI9 were in accordance with the SDs in this region, and exhibited various expressions according to RNA-seq in the spleen and blood (Fig. 6j).

We further investigated the domestication of SVs in bezoars and domesticated goat populations. The top 1% of *F*_ST_ values identified a total of 202 SVs as candidate variants,

63 of which overlapped with the gene bodies (Fig. 6k) and shared 36 genes with those obtained with *ARS1* as the reference genome (Fig. 6l, Supplementary Fig. 19b, and Supplementary Table 11). These 63 genes were associated with immune activity (*MUC6*, *GIMAP8*, *CNTNAP2*, *KCND3*, and *NBEAL1*), stress resistance (*ABCC4*, *PSIP1*, and *TOMM70*), and sperm flagellar function (*SPAG16*), and were classified into three groups. The first group, which included 36 genes, was detected by both *T2T-goat1.0* and *ARS1*; the second, which included 23 genes, were detected in non-PURs of *T2T-goat1.0* but was not found in *ARS1*; and the third, which included 4 genes, was detected only in the PURs of *T2T-goat1.0*. A 62-bp deletion was found in the *MUC6* gene of bezoars on CHI29, which was in accordance with the deletion allele in bezoars compared to goats, as previously reported^2^. Additionally, a deletion (54 bp) was found within *ABCC4* (gene ID: IMCG12g00097) of bezoars on PURs of CHI12 and was confirmed with the IGV^39^ tool (v2.13) (Fig. 6m), whose allele frequencies showed a strong selective signal in domestic goats compared to bezoars. Further examination revealed that the region was annotated with 14 tandem *ABCC4* genes but was not completely assembled in *ARS1*. A duplication (137,204 bp) was found in the myeloid-associated differentiation marker (*MYADM*, IMCG21g00085) of bezoars on the PURs of CHI21 (Supplementary Fig. 20b), which involved 17 *MYADM* in neighboring regions, and *MYADM* was shown to be a host factor essential for parechovirus entry and infection^40^.

### Novel selective signatures underlying the genetic improvement of cashmere traits

To decipher the genetic basis of the cashmere trait, we conducted a selection analysis of the cross-population composite likelihood ratio (XP-CLR) between the cashmere and noncashmere goat populations. Using the top 1% of the XP-CLR score values, we identified a total of 2692 selective signals, spanning 109.22 Mb and covering 1440 genes (Fig. 7a and Supplementary Table 12). In addition to the previously reported cashmere-related *FGF5* on CHI6 and *EDA2R* on the X chromosome previously reported^22^, we have identified selective signals associated with cashmere traits, such as the *KRT* genes on both CHI5 and CHI19, and the *FGF* genes on both CHI5 and CHI20. We also observed selective genes associated with coat color, such as *TYRP1* on CHI8, *KITLG* on CHI5 and *SLC24A4* on CHI21. Compared to those in the *ARS1* reference genome (Supplementary Fig. 21a), we identified 74.36-Mb selective regions uniquely in *T2T-goat1.0* (Fig. 7b). Furthermore, we discovered 274 selective signals located in the PURs, covering 72 genes. A strong signal in the PURs of CHI12 (CHI12: 14.88 Mb – 16.58 Mb), which was supported by the π ratio estimate between the non-cashmere and cashmere goat populations (Fig. 7c), referred to the selection of *ABCC4* genes. The collinearity results indicated incomplete assembly of this region in *ARS1* (Fig. 7d), where 14 tandemly repeated *ABCC4* genes enriched with SDs were assembled in *T2T-goat1.0* (Fig. 7e). We also revealed significant differences in allele frequencies between cashmere and noncashmere populations in the intronic region of one *ABCC4* gene (IMCGChr12g00092, Supplementary Fig. 22a). The *ABCC4* gene is responsible for transporting a range of endogenous molecules and cellular detoxification and is deeply involved in keratinocyte differentiation and stratum corneum keratinization^41^.

**Fig. 7.**
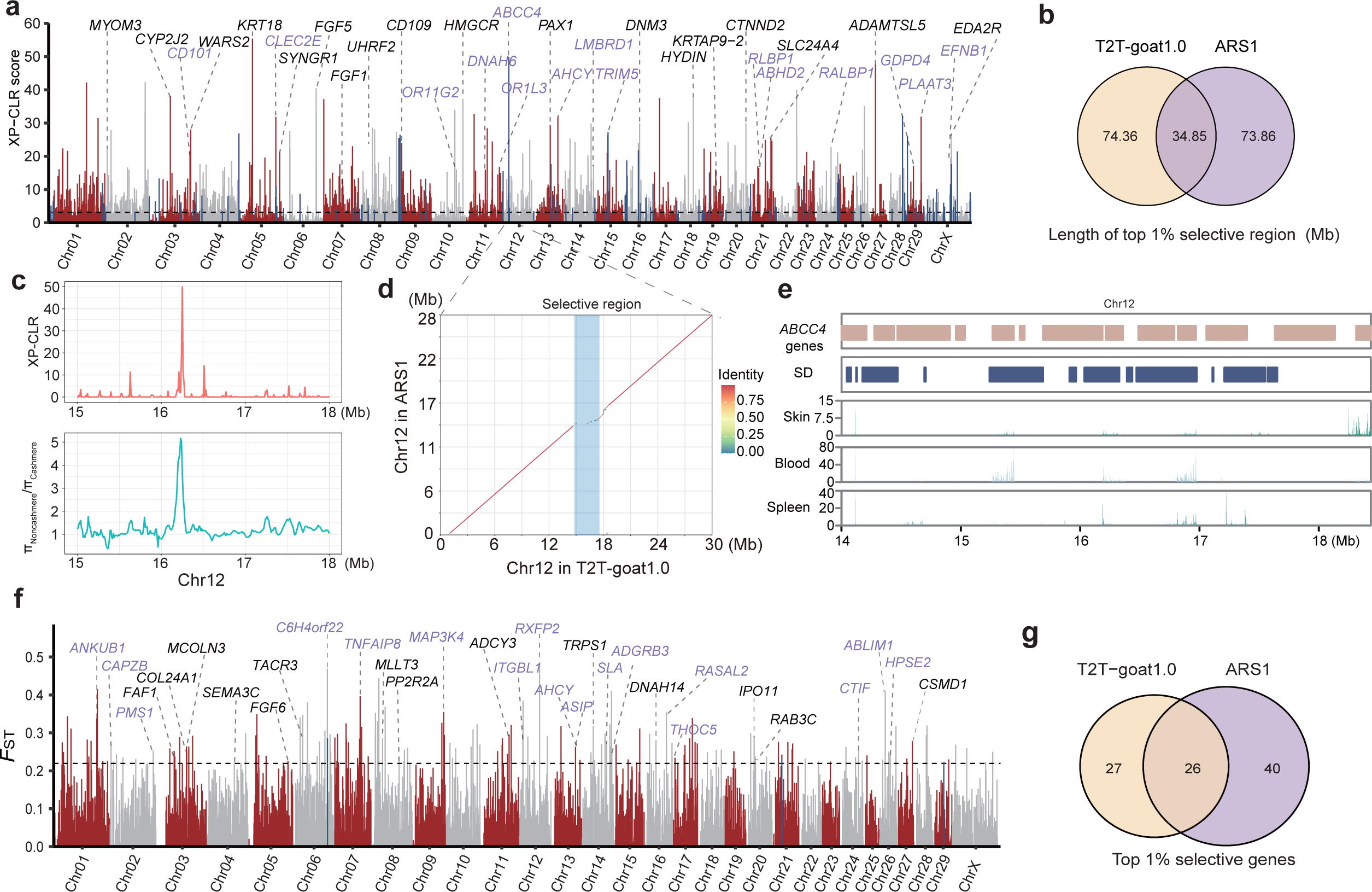
Selection signatures for cashmere traits. **a,** Selective signals based on SNPs and top 1% *XP-CLR* scores for cashmere trait. The selected gene symbols are shown in the same colors to that of Fig. 6e. **b,** Venn plot of selective regions based on top 1% *XP-CLR* scores for the cashmere trait between *T2T-goat1.0* and *ARS1* as references. **c,** Selective region with tandem *ABCC4* genes based on the *XP-CLR* score and π ratio. **d,** Collinearity of the region with tandem *ABCC4* genes on chromosome 12 between *T2T-goat1.0* and *ARS1*. The selective region highlighted in blue bar is exactly located inside the region where tandem *ABCC4* genes are not assembled correctly in *ARS1*. **e,** *ABCC4* genes are in accordance with SDs and showed their expression in skin, blood, and spleen tissues. **f,** Selective signals based on SVs and top 1% *F*_ST_ values between cashmere and noncashmere goats. The selected gene symbols are shown in the same colors to that of Fig. 6e. **g,** Venn plot of selective genes based on top 1% *F*_ST_ scores for the cashmere trait between *T2T-goat1.0* and *ARS1* as references.

In addition to SNPs, 173 selective SVs associated with the cashmere trait were identified based on the top 1% *F*_ST_ values between the cashmere and noncashmere populations (Fig. 7f and Supplementary Table 13). We found that 53 genes related to these SVs were involved in multiple pathways, including those related to hair growth (*FGF6*, *ASIP* and *MCOLN3*), horn formation (*RXFP2*), immune system (*TNFAIP8*, *CSMD1* and *TACR3*), and olfactory system (*OR2T12*). The 26 genes were identified by both the two genome assemblies (Fig. 7g and Supplementary Fig. 21b), and 27 genes were found uniquely by *T2T-goat1.0*. For example, the Agouti-signaling protein (*ASIP*) gene is involved in the regulation of melanogenesis and hair pigment traits in terms of mutation and copy number^42,43^, and was found to be associated with a duplication of 162,498 bp on CHI13 in cashmere goats, which also revealed a differentiation in copy number between bezoars and domestic goats (Supplementary Fig. 22b). Taken together, our selection tests revealed quite a few novel genes and variants associated with cashmere formation during the selection of cashmere traits, and demonstrated the ability of *T2T-goat1.0* to detect selective signals and candidate genes associated with phenotypic traits in goats.

## Discussion

To date, *T2T-goat1.0* is one of several complete seamless mammal genomes, similar to those of human^10,11^, gerbil^44^ and cattle^45^. With the advancement of *T2T-goat1.0*, a black box corresponding to highly repetitive regions was uncovered, especially for centromeric regions. *T2T-goat1.0* revealed three types of centromeric satellite units in goats, with lengths of 816 bp, 702 bp and 22 bp; these findings are supported by the previous studies on centromeric satellite units of 816 bp^25^ and 22 bp^46^. Satellite sequences of the Bovidae family are associated with chromosomal evolution^47^, demonstrating the phylogenetic conversation; however, in mammals, the centromeric satellite repeat unit varies in length and harbors conserved bases in the evolutionarily conserved domain^48^. This phylogenetic conservation could be confirmed by the good alignment of sheep SatI and SatII against *T2T-goat1.0* and the high similarity of SatIII with 83% of the homologs between sheep and goats^46^. Without the isolation and enrichment of satellite DNA by the methods of density gradient centrifugation or restriction enzyme digestion^47^, the T2T genome assembly provided a global view of satellite sequences across each chromosome, and even revealed two variants within SatC of 22 bp (Fig. 3a). The conservation of centromeric satellites could be further investigated on acrocentric chromosomes in the T2T genome assemblies of Bovidae species, for example, cattle^49^, sheep^50^, and deer^51^.

Due to the high repetitiveness and homologous regions with the X chromosome, the complete assembly of the Y chromosome is extremely challenging, while *T2T-goat1.0* successfully resolved the homologous regions with the X chromosome and uncovered 26 gene copies of *TSPY* and 8 copies of *HSFY*. An increase in the number of copies of the testis-specific *TSPY* gene was also reported on the Y chromosomes of human (45 copies)^19^ and cattle (94 copies)^52^. This dosage effect addresses spermatogenesis and is related to the regulation of spermatogenic efficiency^19^. The copy number of *TSPY* reportedly varies significantly among various populations^52^. *HSFY* is believed to be expressed predominantly in the testis and to regulate spermatogenesis by mediating the expression of heat shock proteins (HSPs)^53^. The number of *HSFY* copies varies among the species, with only 2 copies in human^19^, 8 copies in feline^54^, 6 copies in sheep^55^, and 197 copies in cattle^53^. No other multicopy protein-coding gene families were identified in *T2T-goat1.0*. The Y chromosome of goats features a unique centromeric repeat, CenY, which was not found on the other chromosomes with centromeric SatI-SatIII. A unique centromeric repeat unit on the Y chromosome was also found in the other animals; for example, a 1747-bp repeat unit was found in the Mongolian gerbil^44^.

The *T2T-goat1.0* assembly enables more capacity to search for additional selective signatures. First, our assembly revealed 486 NAGs that were not found in *ARS1*. Similarly, T2T genome assemblies added 227 NAGs to the previous human genome reference *CHM13*^10^, and 246 NAGs in maize^14^, 314 NAGs in rice^56^, and 316 NAGs in soybean. Another key point for an addition of selective signatures is that the *T2T-goat1.0* significantly improved repetitive sequences in the goat genome assembly, which facilitated the accurate assembly of tandemly repeated genes under newly identified selection signals, such as *NKG2D* and *ABCC4*. These regions are valuable for selection in terms of both copy number variations and mutations in one of the tandem genes. The strategy of multiple gene copies might be associated with the selection of copy number variations, such as *KIT*^3^ and *ASIP*^57^ for coat color in goats. Remarkable contraction of the *ABCC4* genes was found in domestic goats compared to bezoars^58^, and it was believed that the *ABCC4* genes were involved in the immune system. We confirmed the selection of *ABCC4* (IMCG12g00097) gene during domestication, which showed a new deletion in bezoars. Furthermore, we found that another *ABCC4* gene (IMCG12g00092) was under selection for the cashmere trait, which proved the occurrence of multiple selections in the region with tandemly duplicated *ABCC4* genes, which function in different tissues. Together with other ABC transporters, *ABCC4* genes might be potentially important regulators of epithelial biology in human hair follicles^59^. These genes are under selection for cashmere traits in goats and deserve further investigation. *FGF5* on CHI6 and *EDA2R* on CHIX were previously shown to be associated with cashmere traits^22^, suggesting the reliability of our selection results (Fig. 7a). Additionally, we identified several selective genes that are significantly associated with cashmere traits; for example, multiple *KRT* genes under the strongest selection signal on CHI5 were found (Fig. 7a). Keratin proteins, as a super gene family, are regarded as markers of the secondary hair follicle cycle in cashmere goats^60^. Multiple copies of *KRT* genes were shown to cluster on CHI5, which had the strongest selective signals. We showed strong evidence that these regions consisted of multiple copies of genes under selection in goat genomes, including those with *ABCC4*, *KRT*, *NKG2D*, and *MYADM*. Thus, *T2T-goat1.0* enabled the accurate discovery of these regions and their variants.

In conclusion, the *T2T-goat1.0* assembly enabled a comprehensive evaluation of complex regions with repetitive sequences, improved the alignment of short and long reads, increased the variant calling rate, and identified additional novel selective signatures previously unreported. Population genetics analyses benefited from not only more SNPs in PURs for selective signals but also accurately assembled regions with tandemly duplicated genes.

## Methods

### Sample collection and sequencing

A 4-month-old buck of the Inner Mongolia cashmere goat from the National Preservation Farm for Arbas Cashmere Goat (Ordos, Inner Mongolia, China) was selected for DNA sequencing, RNA-seq and Iso-seq in the T2T genome assembly and subsequent analyses (Fig. 1a). The parents with complete breeding records were also sampled for the assistance of Y-chromosome assembly (Fig. 1a). Additionally, 14 tissues (Supplementary Table 1) from Inner Mongolia cashmere goats on the same farm were collected for RNA-seq and gene annotation.

Five representative goat breeds with specialized traits, i.e., Liaoning cashmere goats with high wool production, Zhongwei goats with white fur and good leather, Jining gray goats with high fertility, Boer goats with high-quality meat, and Tibetan goats with good adaptation to plateaus, were selected for genome sequencing to produce the PacBio long reads (Supplementary Methods), which were used to call SVs (Supplementary Table S1 and Supplementary Fig. 16a).

Blood from the 4-month-old buck and its parents was collected in 10-ml BD Vacutainer blood collection tubes (Cat# 368589, Becton Dickinson, NY, USA). The tubes were placed on dry ice, transferred to the laboratory, and stored at −80°C until DNA extraction. High-molecular-weight genomic DNA was extracted based on the CTAB method, and purified with a Blood & Cell Culture DNA Kit (Cat# 13343, Qiagen, Beijing, China) following the manufacture’s protocols. Library construction and sequencing for PacBio (20-kb libraries, Pacific Biosciences, CA, USA), ultralong ONT (200-kb libraries, Oxford Nanopore Technologies, Oxford, UK), MGI (MGI Tech, Shenzhen, China), Hi-C and Bionano optical maps were all performed at the Grandomics Ltd. (Wuhan, China), and are described in the Supplementary Methods.

All animal procedures were performed in accordance with the guidelines for animal experiments approved by the China Agricultural University Institutional Animal Care and Use Committee (CAU20160628-2).

### Bulk RNA-seq and Iso-seq

Total RNA was extracted from blood using TRIzol reagent in an RNAprep Pure Tissue Kit (Cat# 4992236, TIANGEN Biotech, Beijing, China). After the RNA purity and concentration were determined with Nanodrop (Thermo Fisher Scientific, Waltham, USA) and Qubit (Thermo Fisher Scientific, Waltham, USA), high-quality RNA samples (RIN>8, OD260/OD280=1.8∼2.2, OD260/OD230>2.0) were used for cDNA synthesis in bulk RNA-seq and Iso-seq. Sequencing libraries for Iso-Seq were constructed using the SMRTbell Template Prep Kit 2.0 (Pacific Biosciences, CA, USA), and sequenced on the PacBio Sequel II platform using the Sequel Binding Kit 3.0 (Pacific Biosciences). For RNA-seq, fragmentation and cDNA synthesis were performed using the MGIEasy RNA Library Prep Kit V3.1 (Cat# 1000005276, MGI), and circularization of double-stranded cDNAs was achieved using the splint oligos in the MGIEasy Circularization Kit (Cat# 1000005260, MGI). Single-stranded circlular DNA (ssCir DNA) was used as the final library, and the qualified libraries were sequenced on the DNBSEQ-T7RS platform (MGI). The other 14 tissues (Supplementary Table 1) were also subjected to RNA extraction and Illumina sequencing (details in the Supplementary Methods).

### Fluorescence in situ hybridization (FISH)

The sequences of 45S rDNA, 1474-bp CenY, 702-bp SatII, and 22-bp Sat III were synthesized as probes and labeled with Dig-dUTP or Bio-dUTP (Roche Diagnostics, Basel, Switzerland) using the Biotin and Dig Nick Translation Mix (Roche, Mannheim, Germany). Chromosome preparations were made from fibroblast cell culture as described previously^61^. Slides with cell suspensions at metaphase were hybridized with hybridization mix containing probes and washed for imaging. The hybridization signals were detected based on Alexa Fluor 488 streptavidin (Thermo Fisher Scientific, Waltham, MA, USA) for biotin-labeled probes and rhodamine-conjugated anti-digoxigenin (Roche Diagnostics, Basel, Switzerland) for digoxigenin-labeled probes. Chromosomes were counterstained with DAPI in a mounting medium (Vector Laboratories, Odessa, Florida, USA). Chromosomes in FISH were observed in an Olympus BX63 fluorescence microscope equipped with an Olympus DP80 CCD camera (Olympus, Tokyo, Japan), and images were acquired using cellSens Dimension 1.9 software (Olympus, Tokyo, Japan).

### Draft assembly

For the 4-month-old buck, we generated 141.03 Gb of PacBio HiFi (49.3× coverage) with an N50 of 18.95 kb (Supplementary Table 1), which was used to construct the draft assembly (GV1) using Hifiasm^14^ (v0.16.1-r375) with default parameters. The nonnuclear genome sequences were removed based on the alignment results against the NCBI nucleotide nonredundant database using BLASTN^62^ (v2.10.0) with the parameter of “-evalue 1e-5”. All contigs were aligned to the goat reference genome assembly *ARS1* (GCF_001704415.1) to evaluate the accuracy of GV1 using Minimap2^63^ (v2.26) before further processing.

### Scaffolding with Bionano optical maps

Next, we generated optical maps for scaffolding. Clean Bionano data (molecular length < 150 kb and minSites < 9) of 1209.75 Gb (∼ 362.66× coverage) was used to perform a *de novo* assembly, and generate a consensus map file (.cmap) using the BioNano Solve package (v3.5.1, https://bionano.com/software-downloads/). To generate more continuous consensus genome maps, hybrid scaffolding was conducted between the genome contigs (GV1) and the above genome maps using the “HybridScaffold” module of the Bionano Solve package. The hybrid scaffolds were aligned to the original *in silico* maps from sequence scaffolds using RefAligner (https://bionanogenomics.com/support/software-downloads/), and sequences from GV1 were split at breakpoints (i.e., the conflict positions). The scaffold sequences were produced from the XMAP-formatted alignment and outputted as FASTA files.

Breakpoints generated by the Bionano tools were inspected manually by the Bionano Access software (v1.7; https://bionano.com/software-downloads/), and further verified by the ONT and HiFi data. In summary, ONT and HiFi reads were aligned to the GV1 assembly using Minimap2 with the parameter of “--secondary=no”, and were visualized with the IGV tool (v2.13) to confirm the exact breakpoints. When the ONT and HiFi read alignments supported the original sequences [> 5 reads uniquely aligned and mapping quality (MAPQ) score > 60] where the breakpoints occurred using Bionano tools, the gaps generated by Bionano were abandoned, and the contigs were manually returned. The resulting scaffolded genome (GV2) was subsequently generated to construct the pseudo-chromosome-level genome assembly as follows.

### Pseudochromosome construction

After trimming using fastp (v0.23.1)^64^, high-quality Hi-C reads were aligned to the scaffolded genome assembly (GV2) using Bowtie2^65^ (v2.3.2) with the parameters “-end-to-end --very-sensitive -L 30”. Valid read pairs were generated to identify interactions using HiC-Pro^66^ (v2.8.118). All the sequences were clustered, ordered and oriented onto the 31 pseudochromosomes (29 autosomes and X and Y chromosomes) using the program LACHESIS^67^, with the following parameters: “CLUSTER_MIN_RE_SITES=100, CLUSTER_MAX_LINK_DENSITY=2.5, CLUSTER_NONINFORMATIVE_RATIO = 1.4, ORDER_MIN_N_RES_IN_TRUNK=60, ORDER_MIN_N_RES_IN_SHREDS=60”. The Y chromosome was abandoned due to failed assembly of the homologous region with the X chromosome, and independent assembly was adopted for the Y chromosome with the assistance of the parent’s genome sequences. Placement and orientation errors with obvious discrete chromatin interaction patterns were manually adjusted to assemble the chromosome-level assembly (GV3) of 30 pseudochromosomes (29 autosomes and the X chromosome).

### Gap filling

To fill the gaps, ultralong ONT reads with a length of >100 kb were mapped to all the 30 pseudochromosomes of GV3 using Minimap2 with the option of “-x map-ont”. The ultralong ONT reads that were precisely aligned to the ends of the neighbor contigs connecting the beginning site and ending site of a gap (identity≥95%, coverage≥90%, and QV≥20) were selected as anchor sequences. Short gaps were easily filled by extending the anchor sequences. To fill large gaps, we selected three kinds of ultralong ONT reads based on the read alignment results to conduct local assembly. In addition to the first kind of anchor sequences, we retrieved the second kind of ONT reads, which were roughly aligned to the neighboring regions of the gaps (identity < 95% and coverage < 90%). The reads without any hits against the pseudochromosomes of GV3 were selected as the third kind, as some of them might have originated from one specific gap. The above three kinds of reads were aligned with each other using Minimap2 (v2.23) to produce a library for the overlapping of the pairwise ONT reads. Gaps are mostly located in highly repetitive sequences, such as centromeres, and can result in overlapping errors with a high frequency of *k*-mers. To reduce the false positive overlap caused by repetitive sequences, *k*-mer (*k* = 23) hash tables were calculated based on the MGI short reads using Meryl (v1.4.1)^68^, and rare *k*-mers or low-frequency *k*-mers were determined when the depth was less than the average depth. Subsequently, with the first kind of reads as the starting and ending points, a string graph was built for one specific gap using the Nextgraph module in NextDenovo^69^ (v2.5.2), based on the overlapping ONT reads and their rare *k*-mers. The graph with the longest accumulated length of rare *k*-mers was selected to assemble the sequences to fill the gaps. All ONT reads were aligned to the gap-filled genome to check the coverage and depth of gap regions in IGV (v2.13), and the gap region sequences were adjusted manually with multiple iterations. We subsequently generated the gap-free genome assembly GV4, which covered all the unplaced contigs in GV3.

### Y-chromosome assembly

The Y chromosome was assembled using a haplotype binning strategy and hash tables of *k*-mers from the parents. In brief, *21*-mer libraries were constructed from MGI short reads (∼120× coverage) for the buck individual for the T2T genome assembly and its parents using the Jellyfish^70^ (v2.3.0). Furthermore, the paternal and maternal unique *21*-mers were identified if a *21*-mer was found only in either the father or mother, respectively, and had a sequence depth of >3 and <120 (the average depth across the whole genome) in the T2T buck individual. The ultralong ONT reads were chosen for paternal origination only if more paternal *21*-mers were found than maternal *21*-mers. The paternal ONT reads sequenced from the autosomes were subsequently removed out to retrieve those potentially from the paternal X and Y chromosomes, which were used to construct the assembly graph of the Y chromosome using the Nextgraph module in NextDenovo (v2.5.2). The Y chromosome assembly was further improved by filling gaps and correcting possible assembly errors according to the Y contigs by the trio-binning model of HiFiasm (v0.16.1-r375) based on HiFi reads and *31*-mers from the T2T buck parents. The Y chromosome was combined with all the autosomes and the X chromosome to obtain the genome assembly GV5.

### Telomere assembly

We pooled three types of HiFi reads for independent telomere assembly, namely, those that contained >10 telomere-specific repeats (i.e., AACCCT or AGGGTT), those that were not mapped to the genome assembly GV5, and those that were partially aligned to the 1-Mb chromosomal ends. Telomere assembly was performed using HiFiasm (v0.16.1-r375) with the default parameters, and the contigs with telomeric sequences were aligned to GV5 using Minimap2 to determine their potential chromosomal locations. Telomeres were placed at the 1-Mb ends of each chromosome based on the overlap between contigs and chromosomal ends using RagTag^71^ (v2.1.0, Supplementary Methods). The updated chromosomal ends were manually checked for complete alignments of the HiFi and ONT long reads.

### Genome polishing

To polish the genome assembly GV5, we developed a pipeline (https://github.com/Wuhui2024/CAU-T2T-Goat) that involves long and short reads (Supplementary Figure 23 and Supplementary Methods). First, HiFi reads were mapped to GV5 using Minimap2, and the low-quality regions were then determined based on the HiFi read alignment (those with a MAPQ score ≤1, clipped reads at the both ends, and <3 HiFi aligned reads), while the remaining regions were defined as high-quality regions. The low-quality regions were polished for the first round based on HiFi long reads using Nextpolish2^72^ (v0.1.0), the second round based on ONT long reads and the third round based on MGI short reads using Nextpolish^73^ (v1.4.1). Furthermore, the low- and high-quality regions were both polished based on HiFi long reads using Nextpolish2, before being merged as a whole genome. To avoid the mistakes caused by the merging, the last round of polishing was performed with HiFi reads using Nextpolish2. Next, one additional round of polishing with HiFi reads by Nextpolish2 was conducted to generate the final seamless genome assembly *T2T-goat1.0*.

### Validation of the T2T genome assembly

First, for read coverage analysis, MGI short reads and long reads of ONT and HiFi were aligned to *T2T-goat1.0* using BWA (v0.7.17) and Minimap2, respectively. Genome coverage was subsequently evaluated with a window size of 1 Mb using Bamdst (a BAM depth statistics tool; https://github.com/shiquan/bamdst), and plotted using the KaryoPloteR^74^ package (v1.8.4). For collinearity analysis, the two chromosomes were compared using Minimap2, and genome synteny was visualized using paf2doplot (https://github.com/moold/paf2dotplot) and GenomeSyn^75^. The completeness of the *T2T-goat1.0* assembly was evaluated using BUSCO^23^ (v4.0.5) based on the eukaryota_odb10 database. To evaluate the consensus accuracy, a *k*-mer library (*k* = 21) was built from the MGI short reads using Meryl (v1.4.1), and quality scores per base (i.e., QV) were calculated for the whole genome assembly using Merqury^68^ (v1.3).

### Annotation of repetitive sequences

The methods of *de novo* prediction and homolog-based searching were used to annotate the repetitive sequences including tandem repeats (TRs), transposons, satellites, etc. Four programs, such as GMATA^76^ (v2.2), Tandem Repeats Finder^77^ (TRF, v4.09.1), MITE-Hunter^78^ (v1.0) and RepeatModeler2^79^ (v2.0.4), were applied for *de novo* prediction. The *de novo* repeat libraries were merged into a repetitive sequence database, which was used for homolog-based searching via RepeatMasker^80^ (v4.1.4), with default parameters.

Segmental duplications (SDs) were detected using BISER^81^ (v1.4), and low-quality SDs were filtered when they did not meet the following requirements^82^: 1) Sequence identity >90%, 2) ≤50% gapped sequence in the alignment, 3) Aligned sequence of ≥1 kb, and 4) ≤70% satellite sequence annotated by RepeatMasker.

### Protein-coding gene annotation

Three strategies were used to implement the gene annotation in the *T2T-goat1.0* assembly, namely, the *de novo*, homolog-based, and transcriptome -based approaches. For the transcriptome-based approach, a total of 93 RNA-seq datasets from 40 goat tissues, including 50 ones downloaded from NCBI and 43 newly sequenced ones in this study, were collected (Supplementary Table 1 and Supplementary Table 5), to assemble 145,492 transcripts and obtain 19,437 nonredundant transcripts with Stringtie^83^ (v1.3.4d). Full-length transcripts from Iso-seq were aligned to *T2T-goat1.0* using Minimap2, and redundancies were removed using the “collapse_isoforms_by_sam.py” command of IsoSeq3 (v3.8.2, https://github.com/PacificBiosciences/IsoSeq). Based on the combined nonredundant transcripts from RNA-seq and Iso-seq, gene models were predicted and determined using PASA^84^ software (v2.5.2). For homolog-based prediction, homologous proteins from seven genome assemblies of four closely related species (goat, sheep, human, and mouse) were downloaded from NCBI (Supplementary Table 14), and gene models were determined after alignment to *T2T-goat1.0* using GeMoMa^85^ software. For the *de novo* prediction approach, genes were predicted based on the training set of the above genes in *T2T-goat1.0* with repeats masked by RepeatMasker using AUGUSTUS^86^ (v3.3.1). We integrated all the above predictions of the gene models using EvidenceModeler^87^ (v1.1.1) with default parameters, and the genes with potential transposable elements (TEs) were removed using TransposonPSI (v1.0.0, https://github.com/NBISweden/TransposonPSI). The presence of >700 known genes in previous publications was manually examined to confirm the completeness of the gene annotations in *T2T-goat1.0*. The gene annotations were manually corrected in a modified version of IGV, IGV-GSAman (https://gitee.com/CJchen/IGV-sRNA), based on the transcript evidence.

### Functional annotation of protein-coding genes

To annotate the protein-coding genes, protein sequences were aligned to three databases, Swiss-Prot (http://web.expasy.org/docs/swiss-prot_guideline.html), Kyoto Encyclopedia of Genes and Genomes (KEGG, https://www.genome.jp/kegg/), and the NCBI nonredundant (NR) protein database (ftp:/ftp.ncbi.nih.gov/blast/db/), using BLASTP^62^ (v2.10.0) with the parameters: “-evalue 1e-5 -max_target_seqs 1”. The proteins were also aligned to the Pfam database (https://pfam-legacy.xfam.org/) using InterproScan^88^ (v5.58), and GO terms (http://geneontology.org) were retrieved.

### Centromere identification

A chromatin immunoprecipitation (ChIP) assay was performed to determine the locations of centromeres based on the binding properties of histone-specific antibodies. In brief, 10 ml of fresh blood from the same buck used for the T2T assembly was cross-linked using 1% formaldehyde for 15 minutes at room temperature. The reaction was then quenched by adding glycine at a final concentration of 200 mmol/L. Chromatin was sonicated to obtain soluble sheared chromatin DNA with an average length of 200-500 bp. Afterward, immunoprecipitation was conducted with Phospho-CENP-A (Ser7) antibody (Cat# 2187, Cell Signaling Technology, Beverly, MA, USA). Immunoprecipitated DNA was used to construct sequencing libraries with the NEXTflex ChIP-Seq Library Prep Kit for Illumina sequencing (Cat# NOVA-5143-01; Bioo Scientific, Austin, TX, USA). The final libraries were sequenced on a HiSeq Xten platform with paired-end mode (Illumina, San Diego, CA, USA). After trimming using fastp^64^ (v0.23.1), the clean reads were aligned to *T2T-goat1.0* using Bowtie2 (v2.4.2) with the parameters “--very-sensitive --no-mixed --no-discordant -k 10”. The read depth for ChIP enrichment in 5-kb windows was calculated using BEDTools (v2.30.0) and plotted using the KaryoPloteR package (v1.8.4).

The linguistic sequence complexity was calculated with the program NeSSie^89^ in a window size of 10 kb. The locations of the centromeres were subsequently determined based on the lower sequence complexity due to the enrichment of repetitive sequences^44^. Sequence identity heatmaps for centromeric repeats were generated by StainedGlass^90^ (v0.5), with similar repeats colored according to percentage identity.

### Identification of repetitive sequences inside centromeres

The known 816-bp centromeric satellite DNA sequence (U25964.1) was subjected to BLAST searches against *T2T-goat1.0*, together with ChIP-Seq signals, to preliminarily anchor the centromeric regions. A *k*-mer library was generated based on the above deduced centromere regions using KMC^91^ (v3.1.1) with the parameters “-fm -k151 -ci20 -cs100000”. Based on the centromeric *k*-mers and their frequencies, centromeric repeat units were identified using the program SRF^24^ and were clustered into three groups of minimal satellite repeat units (SatI, SatII and SatIII) based on the sequence identity. The sequence similarity of the goat Y chromosome centromeric repeat unit (CenY, 1473 bp) with the sheep CenY (2516 bp) was determined (Supplementary Methods) using BLASTN (v2.10.0). These centromeric satellite repeats were aligned back to *T2T-goat1.0* to estimate their abundance using BLASTN (v2.10.0). The final centromeric regions were determined based on the distribution of these four types of repeat units.

### Methylation determination by long reads

DNA methylation was calculated based on the HiFi and ONT long reads. HiFi reads were aligned to *T2T-goat1.0* using pbmm2 (v1.13.0) (https://github.com/PacificBiosciences/pbmm2) with the default parameters. The bam file was processed using the “aligned_bam_to_cpg_scores” command of pb-CpG-tools software (v1.0.0) (https://github.com/PacificBiosciences/pb-CpG-tools). The site probabilities of 5-methylcytosine (5mC) methylation were determined and viewed in the IGV program (v2.13). The methylated base (5mC) was also generated from the fast5 format of ultralong ONT reads using Nanopolish^92^ (v0.14.0). Methylation coverage was assessed in a window size of 20 kb using BEDTools^93^ (v2.31.0).

### Structural variation identification

To obtain a high-quality collection of structural variants (SVs), we utilized three tools for the SV calling, namely, pbsv (v2.9.0, https://github.com/PacificBiosciences/pbsv), cuteSV^94^ (v2.0.2), and Sniffles (v2.0.7, https://github.com/fritzsedlazeck/Sniffles). We first aligned the HiFi reads to *T2T-goat1.0* and *ARS1* to generate bam files using Minimap2 and SAMtools^95^ (v1.16). With the program pbsv, we investigated SV signatures based on bam files and called SVs based on PacBio reads for all the five goat samples using the “discover” and “call” modules of pbsv. For the program cuteSV, SVs were called with the parameters of “--max_cluster_bias_INS 1000 --diff_ratio_merging_INS 0.9 --max_cluster_bias_DEL 1000 --diff_ratio_merging_DEL 0.5 --genotype”. For the program Sniffles, we performed SV calling with default parameters. The SVs identified by the three programs were merged using the “merge” command of SURVIVOR^96^ (v1.0.7), with a merging bin of 1000 bp and an SV length >50 bp.

### SNP and SV calling based on short reads

The whole-genome sequence data of caprine genomes, which included 448 domesticated and 68 wild goat individuals, were downloaded from public databases (Supplementary Table 3). First, we performed quality control for the raw sequences using FastQC (v0.11.9; https://www.bioinformatics.babraham.ac.uk/projects/fastqc/) and Trimommatic^97^ (v0.39). Clean short reads were aligned to *T2T-goat1.0* and *ARS1* using BWA^98^ mem (v0.7.17-r1188) with the parameters “-k 23 -M”. The bam files were sorted using SAMtools^95^ (v1.16) and were subsequently used to call SNPs via GATK^99^ (v4.3) as described previously^100^. The raw SNPs were filtered using the VariantFiltration module of GATK with the following parameters: “QD < 2.0 || QUAL < 30.0 || SOR > 3.0 || FS > 60.0 || MQ < 40.0 || MQRankSum < −12.5 || ReadPosRankSum < −8.0”. After removing those with more than 10% missing genotypes and a MAF (minor allele frequency) < 0.05 using VCFtools^95^ (v0.1.16), high-quality biallelic SNPs were retained for subsequent analysis.

Three tools were used to call SVs based on short reads, namely, Delly^101^ (v0.8.7), Manta^102^ (v1.6.0) and Smoove (v0.2.8, https://github.com/brentp/smoove), and the SVs identified by the three programs were merged for all the goat individuals using SURVIVOR (v1.0.7), with a merging bin of 1000 bp and an SV length of >50 bp.

### Population genetics analyses

A phylogenetic tree was constructed from the *p*-distance data using the high-quality autosomal biallelic SNPs and the neighbor-joining (NJ) method in the program PHYLIP^103^(v3.697) and visualized by iTOL^104^ (v6.8.1). LD pruning was performed for independent SNPs using PLINK^105^ (v2.00a3.7) with the following parameters: “--indep-pairwise 50 5 0.2”. PCA was performed on the independent SNPs with the “smartpca” command of the EIGENSOFT^106^ package (v8.0.0). Additionally, population genetic structure was inferred based on the independent SNPs using ADMIXTURE^107^ (v1.3.0). The LD decay (*r*^2^) was measured with a maximum distance of 300 kb using PopLDdecay^108^ (v3.42).

### Selective sweeps during domestication and improvement

To identify selective signals based on both SNPs and SVs, *F*_ST_ values were calculated with 50-kb windows and a 10-kb step between the whole genomes of bezoars and domestic goats using VCFtools (0.1.16). Additionally, we calculated XP-CLR scores using XP-CLR^109^ (v1.1.2) with the parameters “-L 0.95 -P -M 600 --size 50000 --step 10000” and identified potential selective signatures associated with selection for the cashmere traits. Nucleotide diversity (π) was calculated within 100-kb windows using VCFtools (v0.1.16). The top 1% of *F*_ST_ and XP-CLR values were considered as putative selective sweeps associated with domestication and cashmere traits, and the π ratio between the pairwise populations was used to confirm the selective sweeps.

## Acknowledgements

We thank Shaoqing Liu at Inner Mongolia Yiwei White Cashmere Goat Co., Ltd. for helping sampling the T2T buck and the parents, and Xueyan Feng (China Agricultural University) for the help on data analysis. This work was supported by grants from the National Key Research and Development Program-Key Projects (2021YFD1200900 and 2022YFF1003402), the National Natural Science Foundation of China (nos. 31825024, 31661143014, 31972527, 32060742 and 32061133010), and the Second Tibetan Plateau Scientific Expedition and Research Program (STEP; no. 2019QZKK0501).

## Author contributions

M.-H.L. conceived the project. M.-H.L., Z.-H.L. and S.-G.J. supervised the study. H.W. and L.-Y.L. performed genome assembly and data analysis. Y.-H.Z. conducted the experiments on FISH and ChIP-seq, and was involved in generation of the ONT and PacBio data and data analysis. L.-M.Z. performed the telomere analysis, and participated in the discussion of results. J.-H.H. was involved in interpretation and plotting of the results. Z.-H.L., C.-Y.Z., Z.-X.W., and Y.-C.W. performed the RNA-seq sample collection and transcriptome analysis. For PacBio sequencing to call SVs, H.-H.E., Y.Y., H.-D.D., and Z.-Q.Z. collected the blood samples of Zhongwei goat, W.-L.B., D.H., and X.-T. D. collected the blood samples of Liaoning cashmere goat, Y.-L.R., X.-J.W., L.-Y.L. and D.-X.M. collected samples of Jining gray goat, R.D. and L.-Y.L. collected samples of Tibetan goat, and H.-L.C., L.-Y.L. and D.-X.M. collected samples of Boer goat. S.-G.J., H.W., and M.-H.L. wrote the manuscript.

## Data Availability

The genome assembly and raw sequencing data generated in this study, including PacBio HiFi data, ultralong ONT data, MGI data, Iso-seq data and ChIP–seq data, can be achieved from National Genomics Data Center (https://ngdc.cncb.ac.cn/) with BioProject number PRJCA022847.

## Code availability

Custom scripts and codes used in this study are available at GitHub (https://github.com/Wuhui2024/CAU-T2T-Goat). Software and parameters used are stated in the Supplementary Methods with more details.

## References

1. Naderi, S. et al. The goat domestication process inferred from large-scale mitochondrial DNA analysis of wild and domestic individuals. Proc. Natl Acad. Sci. USA 105, 17659–17664 (2008).

2. Zheng, Z. et al. The origin of domestication genes in goats. Sci. Adv. 6, eaaz5216 (2020).

3. Henkel, J. et al. Selection signatures in goats reveal copy number variants underlying breed-defining coat color phenotypes. PLoS Genet. 15, e1008536 (2019).

4. Signer-Hasler, H. et al. Runs of homozygosity in Swiss goats reveal genetic changes associated with domestication and modern selection. Genet. Sel. Evol. 54, 6 (2022).

5. Dong, Y. et al. Sequencing and automated whole-genome optical mapping of the genome of a domestic goat (*Capra hircus*). Nat. Biotechnol. 31, 135–141 (2013).

6. Li, R. et al. A near complete genome for goat genetic and genomic research. Genet. Sel. Evol. 53, 74 (2021).

7. Bickhart, D.M. et al. Single-molecule sequencing and chromatin conformation capture enable *de novo* reference assembly of the domestic goat genome. Nat. Genet. 49, 643–650 (2017).

8. Altemose, N. et al. Complete genomic and epigenetic maps of human centromeres. Science 376, eabl4178 (2022).

9. VarGoats, C. et al. Geographical contrasts of Y-chromosomal haplogroups from wild and domestic goats reveal ancient migrations and recent introgressions. Mol. Ecol. 31, 4364–4380 (2022).

10. Yang, C. et al. The complete and fully-phased diploid genome of a male Han Chinese. Cell Res. 33, 745–761 (2023).

11. Nurk, S. et al. The complete sequence of a human genome. Science 376, 44–53 (2022).

12. Hou, X., Wang, D., Cheng, Z., Wang, Y. & Jiao, Y. A near-complete assembly of an *Arabidopsis thaliana* genome. Mol. Plant 15, 1247–1250 (2022).

13. Deng, Y. et al. A telomere-to-telomere gap-free reference genome of watermelon and its mutation library provide important resources for gene discovery and breeding. Mol. Plant 15, 1268–1284 (2022).

14. Chen, J. et al. A complete telomere-to-telomere assembly of the maize genome. Nat. Genet. 55, 1221–1231 (2023).

15. Wang, L. et al. A telomere-to-telomere gap-free assembly of soybean genome. Mol. Plant, 10.1016/j.molp.2023.08.012 (2023).

16. Li, K. et al. Gapless indica rice genome reveals synergistic contributions of active transposable elements and segmental duplications to rice genome evolution. Mol. Plant 14, 1745–1756 (2021).

17. Huang, Z. et al. Evolutionary analysis of a complete chicken genome. Proc. Natl Acad. Sci. USA 120, e2216641120 (2023).

18. Aganezov, S. et al. A complete reference genome improves analysis of human genetic variation. Science 376, eabl3533 (2022).

19. Rhie, A. et al. The complete sequence of a human Y chromosome. Nature 621, 344–360 (2023).

20. Cechova, M. et al. Dynamic evolution of great ape Y chromosomes. Proc. Natl Acad. Sci. USA 117, 26273–26280 (2020).

21. Guarracino, A. et al. Recombination between heterologous human acrocentric chromosomes. Nature 617, 335–343 (2023).

22. Cai, Y. et al. Ancient genomes reveal the evolutionary history and origin of cashmere-producing goats in China. Mol. Biol. Evol. 37, 2099–2109 (2020).

23. Manni, M., Berkeley, M.R., Seppey, M., Simão, F.A. & Zdobnov, E.M. BUSCO update: novel and streamlined workflows along with broader and deeper phylogenetic coverage for scoring of eukaryotic, prokaryotic, and viral genomes. Mol. Biol. Evol. 38, 4647–4654 (2021).

24. Zhang, Y., Chu, J., Cheng, H. & Li, H. *De novo* reconstruction of satellite repeat units from sequence data. Genome Res. (2023).

25. Nieddu, M. et al. Evolution of satellite DNA sequences in two tribes of Bovidae: A cautionary tale. Genet. Mol. Biol. 38, 513–518 (2015).

26. Burkin, D.J., Broad, T.E. & Jones, C. The chromosomal distribution and organization of sheep satellite I and II centromeric DNA using characterized sheep-hamster somatic cell hybrids. Chromosome Res. 4, 49–55 (1996).

27. Gershman, A. et al. Epigenetic patterns in a complete human genome. Science 376, eabj5089 (2022).

28. Logsdon, G.A. et al. The structure, function and evolution of a complete human chromosome 8. Nature 593, 101–107 (2021).

29. Xiao, C. et al. The assembly of caprine Y chromosome sequence reveals a unique paternal phylogenetic pattern and improves our understanding of the origin of domestic goat. Curr Protoc Bioinformatics 11, 7779–7795 (2021).

30. Shibuya, K. et al. A cluster of 21 keratin-associated protein genes within introns of another gene on human chromosome 21q22.3. Genomics 83, 679–693 (2004).

31. Wang, J. et al. Identification of the caprine keratin-associated protein 20-2 (KAP20-2) gene and its effect on cashmere traits. Genes 8, 328 (2017).

32. Lu, D.D. et al. HAI-1 is required for the novel role of FGFBP1 in maintenance of cell morphology and F-actin rearrangement in human keratinocytes. Hum. Cell 36, 1403–1415 (2023).

33. Ueyama, T. et al. Rac-dependent signaling from keratinocytes promotes differentiation of intradermal white adipocytes. J. Invest. Dermatol. 140, 75–84.e6 (2020).

34. Yu, H. et al. The polymorphism of a novel 30 bp-deletion mutation at *KAP9.2* locus in the cashmere goat. Small Ruminant Res. 80, 111–115 (2008).

35. Jiang, D. et al. Genome array on differentially expressed genes of skin tissue in cashmere goat at early anagen of cashmere growth cycle using DNA microarray. J Integr Agric 13, 2243–2252 (2014).

36. Liu, W. et al. Regular aerobic exercise-ameliorated troponin I carbonylation to mitigate aged rat soleus muscle functional recession. Exp. Physiol. 104, 715–728 (2019).

37. Hu, Z.-L., Park, C.A. & Reecy, J.M. Bringing the Animal QTLdb and CorrDB into the future: Meeting new challenges and providing updated services. Nucleic Acids Res. 50, D956–D961 (2022).

38. Raulet, D.H. Roles of the NKG2D immunoreceptor and its ligands. Nat. Rev. Immunol. 3, 781–790 (2003).

39. Robinson, J.T., et al. Integrative genomics viewer. Nat. Biotechnol. 29, 24–26 (2011).

40. Watanabe, K. et al. Myeloid-associated differentiation marker is an essential host factor for human parechovirus PeV-A3 entry. Nat. Commun. 14, 1817 (2023).

41. Kielar, D. et al. Adenosine triphosphate binding cassette (ABC) transporters are expressed and regulated during terminal keratinocyte differentiation: a potential role for ABCA7 in epidermal lipid reorganization. J. Invest. Dermatol. 121, 465–474 (2003).

42. Adefenwa, M.A. et al. Identification of single nucleotide polymorphisms in the agouti signaling protein (*ASIP*) gene in some goat breeds in tropical and temperate climates. Mol. Biol. Rep. 40, 4447–4457 (2013).

43. Fontanesi, L. et al. Copy number variation and missense mutations of the agouti signaling protein (*ASIP*) gene in goat breeds with different coat colors. Cytogenet. Genome Res. 126, 333–347 (2010).

44. Brekke, T.D. et al. A new chromosome-assigned Mongolian gerbil genome allows characterization of complete centromeres and a fully heterochromatic chromosome. Mol. Biol. Evol. 40(2023).

45. Li, T. et al. De novo genome assembly depicts the immune genomic characteristics of cattle. Nat. Commun. 14, 6601 (2023).

46. Buckland, R.A. Sequence and evolution of related bovine and caprine satellite DNAs: Identification of a short DNA sequence potentially involved in satellite DNA amplification. J. Mol. Biol. 186, 25–30 (1985).

47. Escudeiro, A. et al. Bovine satellite DNAs–a history of the evolution of complexity and its impact in the Bovidae family. Eur Zool J 86, 20–37 (2019).

48. Alkan, C. et al. Genome-wide characterization of centromeric satellites from multiple mammalian genomes. Genome Res. 21, 137–145 (2011).

49. Everts-van der Wind, A., et al. A 1463 gene cattle–human comparative map with anchor points defined by human genome sequence coordinates. Genome Res. 14, 1424-1437 (2004).

50. Jiang, Y. et al. The sheep genome illuminates biology of the rumen and lipid metabolism. Science 344, 1168–1173 (2014).

51. Yin, Y. et al. Molecular mechanisms and topological consequences of drastic chromosomal rearrangements of muntjac deer. Nat. Commun. 12, 6858 (2021).

52. Hamilton, C. et al. Copy number variation of testis-specific protein, Y-encoded (TSPY) in 14 different breeds of cattle (*Bos taurus*). Sex. Dev. 3, 205-213 (2009).

53. Yue, X.-P. et al. Copy number variations of the extensively amplified Y-linked genes, HSFY and ZNF280BY, in cattle and their association with male reproductive traits in Holstein bulls. BMC Genomics 15, 1-12 (2014).

54. Wilkerson, A.J.P. et al. Gene discovery and comparative analysis of X-degenerate genes from the domestic cat Y chromosome. Genomics 92, 329–338 (2008).

55. Li, R. et al. A Hu sheep genome with the first ovine Y chromosome reveal introgression history after sheep domestication. Sci. China Life Sci. 64, 1116–1130 (2021).

56. Shang, L. et al. A complete assembly of the rice Nipponbare reference genome. Mol. Plant 16, 1232–1236 (2023).

57. Norris, B.J. & Whan, V.A. A gene duplication affecting expression of the ovine *ASIP* gene is responsible for white and black sheep. Genome Res. 18, 1282–1293 (2008).

58. Dong, Y. et al. Reference genome of wild goat (*Capra aegagrus*) and sequencing of goat breeds provide insight into genic basis of goat domestication. BMC Genomics 16, 431 (2015).

59. Haslam, I.S. et al. Differential expression and functionality of ATP-binding cassette transporters in the human hair follicle. Br. J. Dermatol. 172, 1562–1572 (2015).

60. Gao, W.Z., Xue, H.L. & Yang, J.C. Proteomics analysis of the secondary hair follicle cycle in Liaoning cashmere goat. Small Ruminant Res. 201, 106408 (2021).

61. Li, X. et al. Genomic analyses of wild argali, domestic sheep, and their hybrids provide insights into chromosome evolution, phenotypic variation, and germplasm innovation. Genome Res. 32, 1669–1684 (2022).

62. Camacho, C., et al. BLAST+: architecture and applications. BMC Bioinformatics 10, 421 (2009).

63. Li, H. Minimap2: pairwise alignment for nucleotide sequences. Bioinformatics 34, 3094–3100 (2018).

64. Chen, S., Zhou, Y., Chen, Y. & Gu, J. fastp: an ultra-fast all-in-one FASTQ preprocessor. Bioinformatics 34, i884–i890 (2018).

65. Langmead, B. & Salzberg, S.L. Fast gapped-read alignment with Bowtie2. Nat. Meth. 9, 357–359 (2012).

66. Servant, N. et al. HiC-Pro: an optimized and flexible pipeline for Hi-C data processing. Genome Biol. 16, 259 (2015).

67. Burton, J.N. et al. Chromosome-scale scaffolding of de novo genome assemblies based on chromatin interactions. Nat. Biotechnol. 31, 1119–1125 (2013).

68. Rhie, A., Walenz, B.P., Koren, S. & Phillippy, A.M. Merqury: reference-free quality, completeness, and phasing assessment for genome assemblies. Genome Biol. 21, 245 (2020).

69. Hu, J. et al. An efficient error correction and accurate assembly tool for noisy long reads. bioRxiv, 2023.03.09.531669 (2023).

70. Marçais, G. & Kingsford, C. A fast, lock-free approach for efficient parallel counting of occurrences of *k*-mers. Bioinformatics 27, 764–770 (2011).

71. Alonge, M. et al. Automated assembly scaffolding using RagTag elevates a new tomato system for high-throughput genome editing. Genome Biol. 23, 258 (2022).

72. Hu, J. et al. NextPolish2: a repeat-aware polishing tool for genomes assembled using HiFi long reads. bioRxiv, 2023.04. 26.538352 (2023).

73. Hu, J., Fan, J., Sun, Z. & Liu, S. NextPolish: a fast and efficient genome polishing tool for long-read assembly. Bioinformatics 36, 2253–2255 (2020).

74. Gel, B. & Serra, E. karyoploteR: an R/Bioconductor package to plot customizable genomes displaying arbitrary data. Bioinformatics 33, 3088–3090 (2017).

75. Zhou, Z. et al. GenomeSyn: a bioinformatics tool for visualizing genome synteny and structural variations. J Genet Genomics 49, 1174–1176 (2022).

76. Wang, X. & Wang, L. GMATA: an integrated software package for genome-scale SSR mining, marker development and viewing. Front. Plant Sci. 7, 1350 (2016).

77. Benson, G. Tandem repeats finder: a program to analyze DNA sequences. Nucleic Acids Res. 27, 573–580 (1999).

78. Han, Y. & Wessler, S.R. MITE-Hunter: a program for discovering miniature inverted-repeat transposable elements from genomic sequences. Nucleic Acids Res. 38, e199 (2010).

79. Flynn, J.M. et al. RepeatModeler2 for automated genomic discovery of transposable element families. Proc. Natl Acad. Sci. USA 117, 9451–9457 (2020).

80. Tarailo-Graovac, M. & Chen, N. Using repeatMasker to identify repetitive elements in genomic sequences. Curr Protoc Bioinformatics 5, 4.10.1–4.10.14 (2009).

81. Išerić, H., Alkan, C., Hach, F. & Numanagić, I. Fast characterization of segmental duplication structure in multiple genome assemblies. Algorithms Mol. Biol. 17, 4 (2022).

82. Vollger, M.R. et al. Segmental duplications and their variation in a complete human genome. Science 376, eabj6965 (2022).

83. Shumate, A., Wong, B., Pertea, G. & Pertea, M. Improved transcriptome assembly using a hybrid of long and short reads with StringTie. PLoS Comput. Biol. 18, e1009730 (2022).

84. Haas, B.J. et al. Improving the A*rabidopsis* genome annotation using maximal transcript alignment assemblies. Nucleic Acids Res. 31, 5654–5666 (2003).

85. Wolf, M. et al. The genome of the pygmy right whale illuminates the evolution of rorquals. BMC Biol. 21, 79 (2023).

86. Stanke, M., Diekhans, M., Baertsch, R. & Haussler, D. Using native and syntenically mapped cDNA alignments to improve *de novo* gene finding. Bioinformatics 24, 637–644 (2008).

87. Haas, B.J. et al. Automated eukaryotic gene structure annotation using EVidenceModeler and the Program to Assemble Spliced Alignments. Genome Biol. 9, R7 (2008).

88. Jones, P. et al. InterProScan 5: genome-scale protein function classification. Bioinformatics 30, 1236–1240 (2014).

89. Berselli, M., Lavezzo, E. & Toppo, S. NeSSie: a tool for the identification of approximate DNA sequence symmetries. Bioinformatics 34, 2503–2505 (2018).

90. Vollger, M.R., Kerpedjiev, P., Phillippy, A.M. & Eichler, E.E. StainedGlass: Interactive visualization of massive tandem repeat structures with identity heatmaps. Bioinformatics 38, 2049–2051 (2022).

91. Kokot, M., Długosz, M. & Deorowicz, S. KMC 3: counting and manipulating *k*-mer statistics. Bioinformatics 33, 2759–2761 (2017).

92. Loman, N.J., Quick, J. & Simpson, J.T. A complete bacterial genome assembled *de novo* using only nanopore sequencing data. Nat. Methods 12, 733–735 (2015).

93. Quinlan, A.R. & Hall, I.M. BEDTools: a flexible suite of utilities for comparing genomic features. Bioinformatics 26, 841–842 (2010).

94. Jiang, T. et al. Long-read-based human genomic structural variation detection with cuteSV. Genome Biol. 21, 189 (2020).

95. Danecek, P. et al. Twelve years of SAMtools and BCFtools. Gigascience 10, giab008 (2021).

96. Jeffares, D.C. et al. Transient structural variations have strong effects on quantitative traits and reproductive isolation in fission yeast. Nat. Commun. 8, 14061 (2017).

97. Bolger, A.M., Lohse, M. & Usadel, B. Trimmomatic: a flexible trimmer for Illumina sequence data. Bioinformatics 30, 2114–2120 (2014).

98. Li, H. & Durbin, R. Fast and accurate long-read alignment with Burrows–Wheeler transform. Bioinformatics 26, 589–595 (2010).

99. McKenna, A. et al. The Genome Analysis Toolkit: a MapReduce framework for analyzing next-generation DNA sequencing data. Genome Res. 20, 1297–1303 (2010).

100. Lv, F.H. et al. Whole-genome resequencing of worldwide wild and domestic sheep elucidates genetic diversity, introgression, and agronomically important loci. Mol. Biol. Evol. 39, msab353 (2022).

101. Rausch, T. et al. DELLY: structural variant discovery by integrated paired-end and split-read analysis. Bioinformatics 28, i333–i339 (2012).

102. Chen, X., et al. Manta: rapid detection of structural variants and indels for germline and cancer sequencing applications. Bioinformatics 32, 1220-1222 (2016).

103. Felsenstein, J. PHYLIP: phylogeny inference package (version 3.2). Cladistics 5, 164–166 (1989).

104. Letunic, I. & Bork, P. Interactive Tree Of Life (iTOL) v5: an online tool for phylogenetic tree display and annotation. Nucleic Acids Res. 49, W293–W296 (2021).

105. Chang, C.C. et al. Second-generation PLINK: rising to the challenge of larger and richer datasets. Gigascience 4, s13742–015-0047-8 (2015).

106. Patterson, N., Price, A.L. & Reich, D. Population structure and eigenanalysis. PLoS Genet. 2, e190 (2006).

107. Alexander, D.H., Novembre, J. & Lange, K. Fast model-based estimation of ancestry in unrelated individuals. Genome Res. 19, 1655–1664 (2009).

108. Zhang, C., Dong, S., Xu, J., He, W. & Yang, T. PopLDdecay: a fast and effective tool for linkage disequilibrium decay analysis based on variant call format files. Bioinformatics 35, 1786–1788 (2019).

109. Chen, H., Patterson, N. & Reich, D. Population differentiation as a test for selective sweeps. Genome Res. 20, 393–402 (2010).

